# Copy number-independent allelic imbalance in mRNA is selected in cancer and has prognostic relevance

**DOI:** 10.1101/2024.09.07.611780

**Authors:** Guillermo Palou-Márquez, Pere Pericot-Masdevall, Fran Supek

## Abstract

Allelic imbalance (AI) in levels of mRNAs in cancer is widely appreciated to result from somatic copy number alterations (CNA) affecting one allele. Apart from CNA, other mechanisms could lead to imbalanced mRNA expression of the alleles, and similarly drive cancer evolution by epistatic interactions with the somatic mutations. By integrating genomic and transcriptomic pan-cancer data, we show that mRNA allelic imbalance favoring the mutant allele in driver genes is subject to positive selection, generating second-hit events often independently of somatic CNA. In some cases, the somatic coding mutations could induce allele-specific expression directly, e.g. with splicing-altering exonic mutations, which can be selected in various cancer genes. However, in the majority of cases, these and related somatic mutation effects (which might in principle alter transcription output via impacting promoters or intragenic enhancers) do not explain the CNA-independent mRNA-level AI, suggesting prevalent epigenetic alterations affecting alleles differently in tumors. Importantly, the mRNA AI events associate with worse overall survival across all cancer types, outperforming various other predictive markers. Our study suggests that mRNA allelic imbalances can occur independently of CNA but similarly function as second-hit events to somatic mutations, driving tumorigenesis, and so represent valuable prognostic biomarkers for cancer patient stratification.

## Introduction

Cancer genomes evolve by accumulation of somatic point mutations, structural variants (which often induce copy number alterations (CNA)), and epigenetic changes. During somatic evolution, healthy cells get transformed because cancer genes, including oncogenes (OGs) and tumor suppressor genes (TSGs), acquire driver point mutations, which largely act by changing the amino acid sequences of the encoded proteins; some structural variants also generate driver changes in a similar manner, by fusing or truncating proteins.

Similarly, cancer genes commonly acquire somatic CNA gains and losses, and epigenetic changes, of which promoter DNA hypermethylation is best studied. These types of alterations can act as tumor drivers by altering gene expression levels – the former, by changing DNA dosage, and the latter, by changing transcriptional output. An important research subject in tumor evolutionary studies is how these various types of driver changes interact with each other. This is epitomized by an observation, well-known from the pre-genomic era, that TSGs bearing damaging germline mutation often require a “second-hit” alteration such as to cause cancer, be it a somatic CNA deletion(1, 2) or an epigenetic change(3). Similarly, TSGs and also OGs bearing damaging somatic mutations are often additionally affected by CNA, acting as second-hit driver events(4, 5).

In diploid genomes such as the human, each gene bears two copies referred to as alleles; somatic driver point mutations typically affect only one allele, with only rare observations of double somatic mutations in the same driver gene(6). The somatic alterations that act via changing gene expression levels – CNA and epigenetic changes – can affect the *mutant* and the *wild-type* OG or TSG alleles differently, constituting an allelic imbalance (AI). The somatic CNA generate an AI at the DNA level, and favor increasing the proportion of the mutant allele of the driver gene by either CNA gains of the mutant or by CNA losses of the *wild-type* allele i.e. a loss-of-heterozygosity (LOH)(5, 7). Effects of these DNA-level AI resulting from CNA propagate to the gene expression, resulting in mRNA-level AI. This was reported for impactful somatic mutations in individual examples of cancer genes (e.g. *PIK3CA*(8)) or cancer types(9–13) (e.g. neuroblastoma(14) [MIM: 256700]), and in two broader studies of thousands of RNA-Seq cancer samples(15, 16). The mRNA-level AI can also be referred to as allele-specific expression (ASE) of the mutant transcripts; importantly here we do not imply that the studied mutations are in themselves necessarily causal to the imbalanced mRNA expression(17).

Apart from the cancer context and the somatic CNAs(4, 15, 18, 19), allelic imbalance of gene expression is a known occurrence in the human genomic studies that consider germline variants(20–23). While the mRNA-level AI can be often caused by *cis*-acting genetic variation – via occurrence of single nucleotide variants (SNVs) that alter gene *cis*-regulatory elements (CREs) such as gene promoter or enhancers(22, 23) or that alter genic regions to disrupt splicing elements therein(24) or introduce stop codons and trigger nonsense-mediated mRNA decay (NMD)(25, 26)– the mRNA AI can also arise by epigenetics. A well-known mechanism generating mRNA allelic imbalances in a defined set of human genes is genomic imprinting(27), in which only a specific parental copy of affected genes is expressed and the other is silenced through epigenetic mechanisms(28). Beyond imprinting and chromosome X inactivation in females, there are reports of mitotically-stable, random monoallelic expression events affecting many genes (reviewed in ref. (29)), associated with a histone mark signature(30), close to topologically associated domains proximal to telomeres(31) or linked with genetic disease penetrance(32). This suggests epigenetic effects may commonly generate mRNA-level AI in human cells, however the prevalence of these epigenetic AIs in cancer, their interactions with driver mutations and resulting impact on tumor evolution were less studied(9, 13, 33, 34).

In this analysis, we compare allelic frequencies of driver somatic mutations between the tumor genome sequencing data (DNA-level, representing somatic CNA effects) and the tumor transcriptome (RNA-level), to dissect the effects of CNA versus other, non-CNA mechanisms causing mRNA allelic imbalance by altering transcription activity or transcript processing(8, 26). A particular hypothesis of interest is whether the non-CNA mechanisms generating mRNA-level AI can modulate selective pressures acting on driver genes, steering cancer evolution, in an analogous way to well-established mechanisms of CNA.

By integrating whole-exome sequencing (WES) and RNA-Seq from 8, 809 tumors(35), spanning 31 different cancer types, we find that non-CNA driven AI of somatic mutations is common in cancer genes, that it is under positive selection, and that it has strong prognostic accuracy of aggressive tumors.

We further suggest that the occurrences of non-CNA AI on coding somatic mutations largely do not result from the mutation itself, even though we note some occurrences of splicing-altering or transcription-activity altering mutations. Rather, the non-CNA AI would result from a separate genetic or more likely epigenetic change, which acts as a second-hit driver event on the cancer gene and manifests as changed DNA methylation and/or chromatin accessibility.

## Methods

### TCGA datasets

We downloaded matched whole-exome sequencing (WES) aligned tumor and normal *bam* files and RNA-Seq tumor raw *fastq* data from the The Cancer Genome Atlas (TCGA consortium)(35), using the Genomic Data Commons (GDC) Data Portal (https://portal.gdc.cancer.gov/), from which 8, 809 samples spanning 31 different cancer types were retained for AI analyses. Tumor purity estimates were also downloaded accordingly with TCGA data(35). Copy number alteration (CNA) gene-level data, for arm-level, was obtained from the Broad Institute’s GDAC Firehose portal (https://gdac.broadinstitute.org/). ATAC-Seq *fastq* data for the analysis of allele-specific chromatin accessibility (ASCA) was also downloaded from TCGA through the GDC Data Portal, from which 381 samples spanning 23 different cancer types were retained. Additionally, DNA methylation data, represented as beta values for 486, 427 CpGs across 8, 689 samples and 32 cancer types, was retrieved using the *TCGAbiolinks* R package(36).

For the non-coding mutations assessment, we downloaded whole-genome sequencing (WGS) files containing somatic variant calls for 777 tumor samples across 22 different cancer types from the Pan-Cancer Analysis of Whole Genomes (PCAWG) dataset (available at https://dcc.icgc.org/releases/PCAWG/Hartwig). These variant calls were processed using the Hartwig analytical pipeline v5 (https://github.com/hartwigmedical/pipeline5)(37). The procedures followed in the analyses were in accordance with the ethical standards of the data providers.

### Sequencing data alignment

RNA-Seq and ATAC-Seq sequencing reads were aligned to the human genome *GRCh38.d1.vd1* using *STAR* v2.5.3a(38). Reads with a minimum mapping quality of 255 were retained and allele-mapping bias was corrected using *WASP*(39). An unbiased removal of duplicate reads was performed using *WASP*-based in-house script (https://github.com/bmvdgeijn/WASP), which discards one of the duplicate reads at random.

### Sequencing data variant calling and annotations

Variant calling was performed using *Strelka2* v2.9.10(40). For WES, we compared normal and tumor *bam* files using *configureStrelkaSomaticWorkflow.py* with flags --exome --tumorBam and --normalBam. For RNA-Seq and ATAC-Seq, we used only tumor bam files, using *configureStrelkaGermlineWorkflow.py with flags --exome --rna --forcedGT*. This last argument forces the calling in the previously called somatic genotypes from WES. SNVs and indels were called separately in all three cases and merged afterwards.

*VCF* files were annotated using ANNOVAR(41) (2020-06-07 version) with GENCODE v26 (ENSEMBL v88) and adding *gnomad211_exome* database to obtain the population minor allele frequencies (MAF) from gnomAD(42). Gene-level expression (as transcripts per million (TPM)) quantification was conducted using *RSEM* v1.3.0(43).

### Data quality control processing

We selected heterozygous synonymous and missense SNVs, as well as nonsense variants, that had a total number of at least 5 RNA-Seq (or ATAC-Seq, depending on the analysis) read counts covering the locus (both *mutated* and *wild-type*), and a “PASS” at the FILTER column obtained from *Strelka2* WES variant calling. Additionally, missense mutations located within genes also bearing nonsense, frameshift indels, or splice site mutations in the same tumor sample were discarded.

### Variant pathogenicity predictions

Variant pathogenicity predictions were obtained using *VARITY*(44, 45), and downloaded from http://varity.varianteffect.org/. Missense variants were assigned an estimated pathogenicity score according to the “*VARITY_ER*” model. Variants mapping to multiple scores were assigned the average of their predicted scores. Missense variants with a predicted *VARITY_ER* < 0.5, were relabeled, along with all synonymous SNVs, as effectively synonymous (ES) variants, while >= 0.5 *VARITY_ER* implies a high amino acid (aa)-impact missense variant (likely pathogenic).

### Functional classification of genes

Data for the classification of individual genes according to their role in cancer was obtained from the Cancer Gene Census (CGC)(46). CGC genes with “Mutation Types” labelled as “N”, “S” and “Mis” were kept, removing mostly fusion and/or translocated genes. The remaining genes were classified according to their annotated role in cancer as TSGs (n = 161), OGs (n = 108), or both TSG/OGs (n = 70). Additionally, the *TP53* gene was treated as a separate group, as there were a high number of variants from that gene. Lists of essential genes were obtained from the CEG2 set determined from CRISPR screens on cell lines(47) and removed from all analysis (n = 626). All remaining genes were classified as passenger genes (n = 15, 258).

Data on the association of genes with specific cancer types was obtained from the MutPanning catalog of significantly mutated driver genes(48). Variants from cancer genes were classified as “cognate” on those cancer types with an existing association in MutPanning at FDR <= 25%, and “noncognate” otherwise. For the selection analysis, we additionally re-classified 16 passenger genes as cancer genes as they were found in the MutPanning catalog.

### Estimating mutant copy number fraction

To estimate mutant copy number (the number of chromosome copies containing the mutation), we applied the formula from Black *et. al.*(16) and McGranahan *et. al.*(49). This incorporates tumor purity, total copy number (derived from ASCAT(50)), and normal copy number:

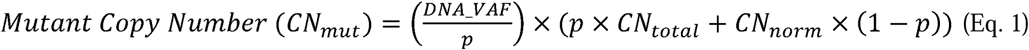

Where 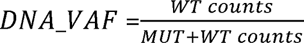, *p* = sample tumor purity, *CN_total_* = total copy number (obtained from ASCAT), and *CN_norm_* = normal copy number (set to 2).

We then divided the mutation copy number by the total tumor copy number at the mutated position, capped at 1, to obtain the mutant copy number fraction:

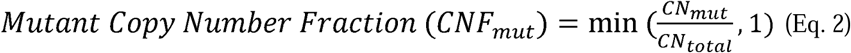

### Allele-specific copy number assignment

We utilized ASCAT(50) major and minor allele copy numbers to assign allele-specific copy numbers to mutant (MUT) and wild-type (WT) alleles based on the previously estimated copy number mutant fraction (*CNF_mut_*):

If *CNF_mut_*< 0.5: ASCAT Major Copy Number → WT allele (*ACN_Wt_*), and ASCAT Minor Copy Number → MUT allele (*ACN_mut_*).

If *CNF_mut_* > 0.5: ASCAT Major Copy Number → MUT allele (*ACN_mut_*), and ASCAT Minor Copy Number → WT allele (*ACN_Wt_*).

Based on these allele-specific assignments, we classified SNV events harboring copy-numbers, depending on the analysis as: (i) copy-number neutral (*CN_neut_*): *ACN_wt_* ≥1 and *ACN_mut_* ≥ 1, (ii) Based on these allele-specific assignments, we classified SNV events harboring copy-numbers, copy-number gain (*CN_gain_*): *ACN_mut_* > 1, (iii) loss-of-heterozygosity (LOH): *ACN_Wt_* ≥ 0; (iv) no LOH: *ACN_Wt_* ≥:1.

### Beta-binomial modeling of allelic imbalance

We developed a framework of beta-binomial models using the *VGAM* R package and ‘*vglm’* function to model allelic imbalance (AI) separately for DNA and RNA data. This approach allows us to quantify overall AI at the mRNA level, copy number alteration-driven AI (CNA-AI), and transcriptional regulatory effects beyond CNA (reg-AI). The general beta-binomial model is formulated as:

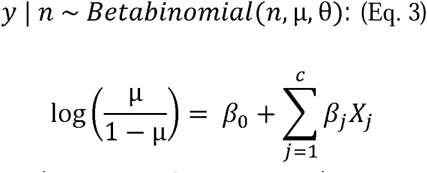

Where *y = MUT* allele counts (number of successes), *n = MUT + wI* allele counts (total trials), µ = mean proportion/success probability or *VAF_pred_*, θ = overdispersion parameter, *β_j_ x_j_* = regression coefficients (*β_j_*) to be estimated from *j* = 1 to *c* covariates (*x_j_*).

Specifically, for **mRNA-AI** we modeled baseline RNA allele counts from RNA-Seq data to capture the total imbalance in the transcriptome. This accounted for potential confounders like tumor purity and overall gene expression level (TPM), including their interaction, to account for the expression contribution of normal cells relative to tumor cells.

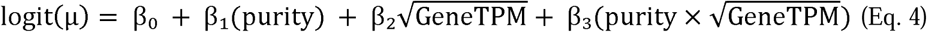

For **CNA-AI**, we modeled DNA allele counts from WES data using two models: a baseline model with sample tumor purity, and a second model that included the expected mutant copy number fraction (*CNF_mut_*) at the DNA level as a covariate (estimated using ASCAT-derived CNA values(50), see section “Estimating mutant copy number fraction”). Note that *CNF_mut_* already incorporates purity in its formula. By comparing these models, we could determine whether observed DNA-level AI was attributable to copy number alterations.

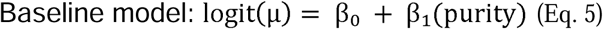

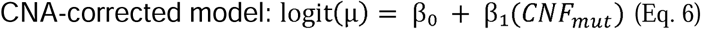

For **reg-AI**, we assessed AI at the mRNA level beyond that attributable to genomic dosage changes. This model used RNA allele counts as the response variable, similar to the overall mRNA-level AI model, but incorporated the expected *CNF_mut_* as an additional covariate to correct for CNA effects.

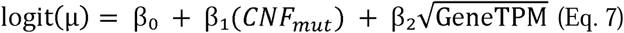

For each somatic heterozygous SNV, *i*, and each AI model separately, *j*, we determined significant AI by testing whether the *VAF_obs_* (observed variant allele frequency) significantly deviated from the *VAF_pred_* (predicted VAF from the model). Specifically, we assessed the probability of observing allele-specific read counts at least as extreme as those observed, given the *VAF_pred_* from the beta-binomial models from above. We used the *’pbetabinom’* function from *VGAM* with the model’s fitted overdispersion parameter to calculate tail probabilities:

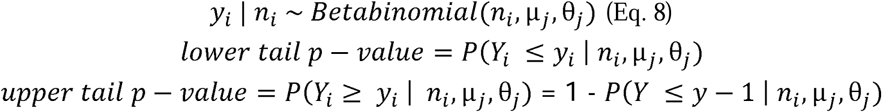

Where *y_i_* = *MUT* allele counts (either RNA or DNA) for an *SNV i, n_i_ = MUT + VT* allele counts (either RNA or DNA) for an *SNV_i_, µ_j_ = VAF_pred_* by the beta-binomial model *j* (either mRNA-AI, CNA-AI or reg-AI), and θ*_j_* = overdispersion parameter of the beta-binomial model *j*.

Given the above tests, SNVs were classified to determine AI status for mRNA-AI, reg-AI, and CNA-AI using its corresponding *j* model test, as follows:

#### Positive AI (pos-AI)

- For mRNA-AI and reg-AI: *RNA_VAF_obs_* is significantly greater than *RNA_VAF_pred_* in their respective RNA models (upper-tail *p* < 0.05) and *DNA_VAF_obs_* is not significantly higher than *DNA_VAF_pred_* in the baseline DNA model (upper-tail *p* ≥ 0.05).
- For CNA-AI: *DNA_VAF_obs_* is significantly greater than *DNA_VAF_pred_* in the baseline DNA model (upper-tail *p* < 0.05), and this significance is absent in the CNA-corrected DNA model (upper-tail *p* ≥ 0.05).

#### Negative AI (neg-AI)

- For mRNA-AI and reg-AI: *RNA_VAF_obs_* is significantly lower than *RNA_VAF_pred_* in their respective RNA models (lower-tail *p* < 0.05) and *DNA_VAF_obs_* is not significantly lower than *DNA_VAF_pred_* in the baseline DNA model (lower-tail *p* ≥ 0.05).
- For CNA-AI: *DNA_VAF_obs_* is significantly lower than *DNA_VAF_pred_* in the baseline DNA model (lower-tail *p* < 0.05), and this significance is absent in the CNA-corrected DNA model (lower-tail *p* ≥ 0.05).

#### No AI

- For mRNA-AI and reg-AI: No significant allelic imbalance is observed in their respective RNA models (*p* ≥ 0.05 for both tails).
- For CNA-AI: No significant allelic imbalance is observed in the baseline DNA model (*p* ≥ 0.05 for both tails).

To clarify these classifications, pos or neg mRNA-AI indicates an overall RNA imbalance (e.g., *mutant* allele bias for pos-AI, *wild-type* bias for neg-AI), assessed using a baseline RNA model. Pos or neg reg-AI indicates an RNA imbalance that persists even after correcting for CNA effects, as it is assessed using a CNA-corrected RNA model. Pos or neg CNA-AI signifies that an initial DNA imbalance observed in the baseline DNA model is subsequently resolved (i.e., disappears) after CNA correction, implying the imbalance was primarily driven by copy number changes.

### AI-dN/dES test for selection

We developed two complementary approaches to test for positive selection on mRNA allelic imbalance: (i) gene category-level analysis, aggregating somatic SNVs across functionally related genes (cognate and noncognate TSGs and OGs; *TP53* separately; passenger genes as controls), and (ii) individual gene-level analysis, examining selection within specific genes (e.g. *KRAS*).

For gene category-level testing, we performed enrichment analyses using Fisher’s exact test to calculate odds ratios (ORs) comparing the prevalence of positive allelic imbalance (reg-AI or CNA-AI, analyzed separately) versus no AI. We contrasted high aa-impact missense mutations against a baseline of ES mutations — conditional “dN/dES” framework — excluding nonsense mutations from the analysis. This approach tests whether high aa-impact missense mutations exhibit preferential expression of the mutant allele compared to effectively-synonymous mutations.

To identify individual cancer driver genes subject to selection pressure on mRNA allelic imbalance, we developed AI-dN/dES, a dN/dS-type test. This approach adapts our previously established pan-cancer beta-binomial framework for detecting significant AI at the single-SNV level (see section “Beta-binomial modeling of allelic imbalance”), applying it here on a per-gene basis. The AI-dN/dES methodology employs gene-specific regression models for both mRNA-AI and reg-AI. The models are defined as follows:

mRNA-AI:

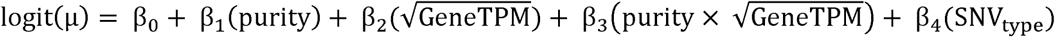

reg-AI:

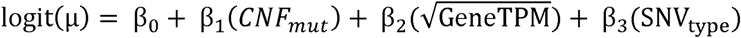

Where µ represents the *VAF_pred_* by the beta-binomial model *j* (either mRNA-AI or reg-AI), purity is the tumor sample purity, GeneTPM is the gene expression level in TPM, *CNF_mut_* is the mutant allele’s copy number fraction (used in the CNA-corrected model), and *SNV_type_* is a categorical covariate representing the functional impact of the SNV.

The model incorporates an *SNV_type_* covariate, and its regression coefficient measures the gene-specific differential AI between missense and a neutral baseline of effectively synonymous mutations. Nonsense mutations were excluded due to potential confounding effects from NMD. A positive, statistically significant effect size (by upper tail *p*-value) indicates that high aa-impact missense mutations drive greater enrichment of the *mutant* allele in RNA than the neutral baseline of ES variants. This signature provides direct evidence that AI itself is under positive selection in the identified driver genes.

For this analysis, we focused on known cancer driver genes from CGC to reduce multiple-testing burdens (see section “Functional classification of genes”). We also included putative passenger genes identified by MutPanning(48) that were not listed in CGC, leading to a total of 356 genes for testing (108 OGs, 162 TSGs, 16 found in MutPanning only and 70 both TSG/OG).

Tested genes were required to have at least 10 somatic SNVs in total, with >3 ES variants and >3 high aa-impact missense mutations. Following the regression analyses, outlier genes were further excluded based on the Z-score of the *SNV_type_* coefficient (|Z-score| > 15).

Following these AI-dN/dES tests, genes with a FDR < 10% (calculated separately for reg-AI and mRNA-AI results) were considered to exhibit significant positive selection on RNA allelic imbalance. Putative selected genes (FDR < 25%) were further considered and included in our downstream survival analysis (see section below).

### Survival analyses

To perform survival analyses, we downloaded clinical data for TCGA from GDC (https://portal.gdc.cancer.gov/). The analyses were conducted using the *survival* package for R(51, 52) and the *survminer* package(53) for Kaplan-Meier (KM) curves with *p*-values obtained through log-rank tests. Alive patients were censored at the end of their follow-up.

Survival outcomes were evaluated for patients with at least one somatic high aa-impact missense mutations in any of the genes identified as significantly positively selected in our AI-dN/dES test. This was done separately for genes selected based on mRNA-AI (35 genes), reg-AI (22 genes). For this survival analysis, the FDR threshold for gene selection was relaxed to 25%, up from the prior 10% threshold, to increase sample and variant retention and boost statistical power. ES variants were excluded. *TP53* was omitted here to avoid bias from its exceptionally high mutation frequency. Within these selected gene sets, patients whose tumors exhibited positive AI were compared against those with no significant AI in the same genes.

To quantify the extent of AI for each tumor, we first calculated the Δ*RNA_VAF* for each qualifying SNV as the difference between the *RNA_VAF_obs_* and the *RNA_VAF_pred_* by our beta-binomial models. The mean of these Δ*RNA_VAF* values across all considered SNVs within a given tumor was then used as a continuous AI score for that sample. Tumor samples were subsequently classified into four distinct categories based on this continuous AI score: neutral/reference (mean Δ*RNA_VAF* score in [-0.1, 0.1]); and positive AI groups based on quantiles of the positive mean Δ*RNA_VAF* scores (mRNA-AI: (0.1, 21], (0.21, 0.37] and (0.37-0.7]; reg-AI: (0.1, 0.16], (0.16, 0.26], and (0.26, 0.68]).

To account for multiple covariates, Cox proportional hazards models were employed at various levels (pan-cancer, individual cancer types, and individual genes) using the *survival* package in R. The models were applied across 31 distinct cancer types and also focused separately on each positively selected gene to assess the individual impacts. CNA gene-level data was obtained for all cancer types separately, downloaded from the Broad Institute’s GDAC Firehose portal (http://firebrowse.org/). CNA burden was measured as the total number of CNA events in genes per sample (discretized to quartiles). Genetic intratumor heterogeneity data (estimated as copy-number heterogeneity, CNH) was obtained from a previous study(54), and was discretized to quartiles. Sex of the patient, tumor sample purity, tumor stage, and age of the patient (discretized as quintiles) were also included in the model as covariates. Results were plotted using the *forestmodel* package for R(51).

Cancer types with sample sizes below 50, or with fewer than 5 samples with available survival information were excluded, leaving 15 cancer types for the Cox regression. Furthermore, individual models were discarded if estimated hazard ratios (HRs) were excessively large (HR>10) or if CIs were “Infinite” or had an upper bound > 10, as these often indicate modelling difficulties. When analysis was stratified by cancer type, the “sex” variable was removed for sex-specific cancers (BRCA [MIM: 114480], UCEC [MIM: 608089], CESC [MIM: 603956], UCS, TGCT [MIM: 273300], OV [MIM: 167000], PRAD [MIM: 176807]); the “stage” variable was omitted if missing in over 90% of samples; the “tumor purity” variable was excluded if it was not available (LAML [MIM: 601626], THYM, and DLBC).

### Mechanisms underlying regulatory allelic imbalance in cancer driver genes

A full description of the analytical steps and data source and preprocessing used to elucidate the causal mechanisms of regulatory allelic imbalance (reg-AI) in our cancer cohort is provided in the Supplemental Material and Methods. The relevant subsections are:

1. Allele-Specific Chromatin Accessibility (ASCA) analysis
2. Gene-level DNA-methylation status
3. Variant effect predictions on chromatin states (Sei)
4. Variant effect predictions on transcription Initiation (Puffin-D)
5. Variant effect predictions on splicing and enrichment analysis
6. Classification of nonsense mutations into NMD-triggering and NMD-evading

## Results

### Modelling allelic imbalance on somatic driver mutations in cancer

In this study, we conceptualize that mechanisms generating AI on cancer driver mutations can occur at two stages: i) somatic CNA events that alter dosage at the DNA level (henceforth, CNA-AI), and/or ii) other mechanisms altering allelic expression at the mRNA level independently of DNA dosage; We attribute this latter, non-CNA AI to epigenetic variation, or to genetic variation impacting regulatory regions, thus influencing transcriptional regulation. Hence, we refer to it as reg-AI (regulatory AI).

The observed differential abundance between allelic transcripts at the mRNA level (henceforth, mRNA-AI), is therefore the net composite of both CNA-AI and reg-AI. Thus, there are different paths by which allelic imbalance could occur, with respect to a somatic mutation (simple schematic in Fig. 1A and detailed schematic in Figure S1), involving imbalance of the mutant allele only at DNA dosage level via CNA-AI, only at mRNA dosage level via reg-AI, or a mixture of both (of note, we eschew the term “allele-specific expression”, because of an interpretation(17) that the imbalanced genetic variants were directly causal to the mRNA imbalance; we do not imply this in our study of somatic driver mutations).

**Figure 1.**
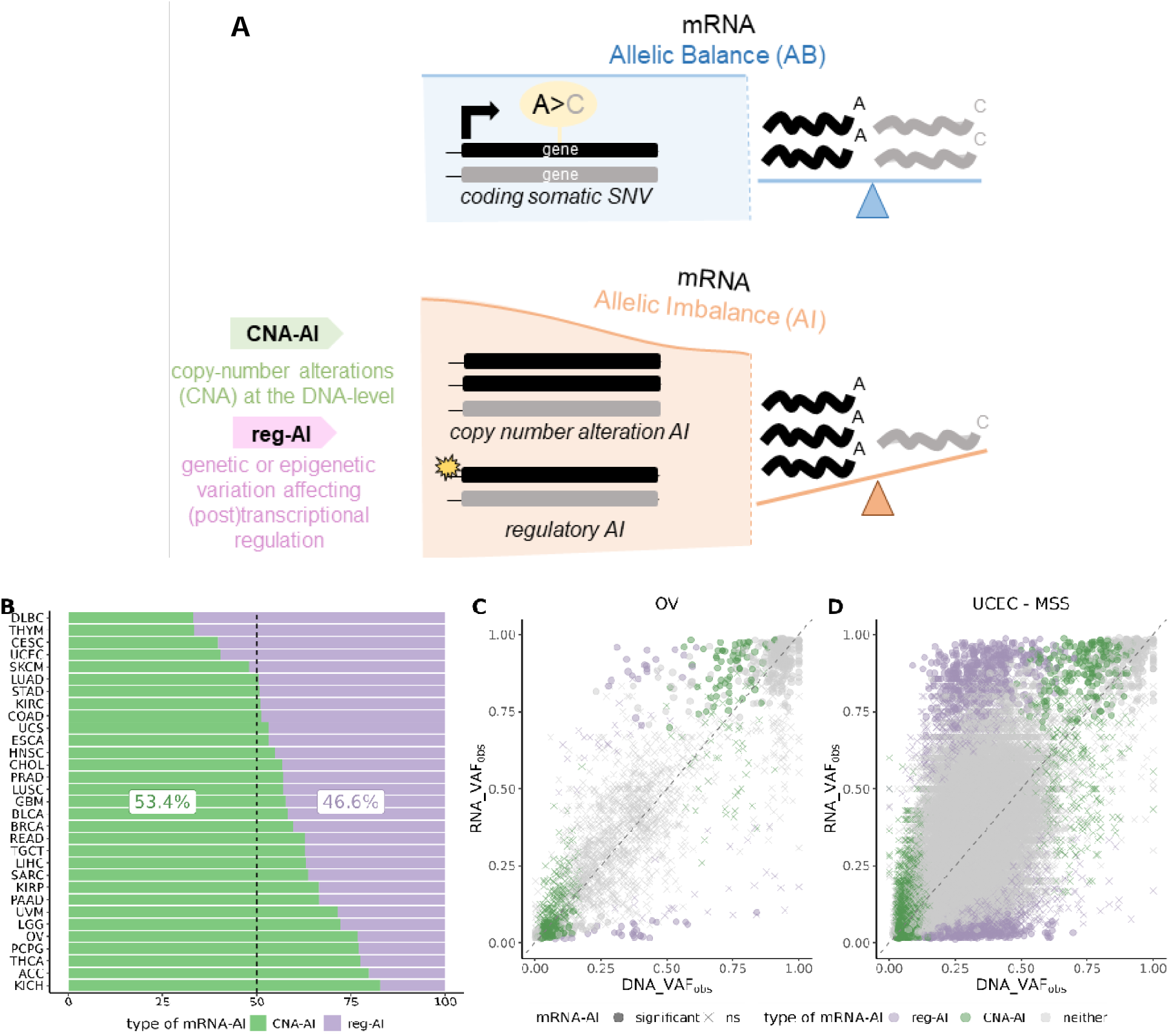
Determinants of allelic imbalance (AI) in cancer genomes. **A**, Illustration of two distinct paths through which allelic imbalance (AI) may favor the mRNA expression of the mutant allele following an somatic coding SNV event: (1) by copy number alterations (CNA) alone or (2) by genetic or epigenetic variations. We attribute this latter, non-CNA AI, to epigenetic changes and/or to genetic variations that influence *cis*-regulatory elements, thereby affecting transcriptional regulation. We collectively refer to these non-CNA mechanisms as regulatory AI (reg-AI). **B**, Proportion of exome- wide SNVs exhibiting significant reg-AI or CNA-AI within the subset of SNVs that demonstrate significant overall mRNA-AI (and additionally significant reg-AI or CNA-AI), stratified by cancer type across the TCGA cohort. It illustrates that AI caused by non-CNA mechanisms has a major impact on overall AI and is pervasive in cancer genomes (46.6% of all SNVs with significant mRNA-AI). **C-D**, Representative examples illustrating observed RNA variant allele frequency () versus distributions for SNVs in two distinct cancer types. SNVs with significant mRNA-AI are highlighted and further categorized by their association with either reg-AI or CNA-AI. In ovarian serou cystadenocarcinoma (OV, **C** panel), the majority of SNVs with significant mRNA-AI are attributable to CNA-AI, as evidenced by the strong correlation between and for these variants. In contrast, for the microsatellite stable subtype of uterine corpus endometrial carcinoma (UCEC-MSS, **D** panel), a larger proportion of SNVs with significant mRNA-AI are driven by reg-AI. These reg-AI associated SNVs typically exhibit *RNA_VAF_obs_* that are significantly higher or lower than their corresponding *DNA_VAF_obs_*, consistent with allelic imbalance at the mRNA level not being explained by DNA dosage alterations (CNA). List of cancer types acronyms can be found in Table S1. TSS, transcription start site; eQTL, expression quantitative trait loci.

Our approach consisted in separating the known effect of CNA-AI from the other effects on mRNA allelic expression imbalance, considering each somatic SNV mutation in cancer exomes, to understand the implications of reg-AI on cancer evolution via positive selection affecting driver genes. Prior studies of allelic imbalances in cancer compared transcriptomes of healthy versus cancerous tissues, quantifying AI of presumably neutral germline variants used as markers of allele-specific expression(4, 10, 15, 18, 19). In contrast, our work considers somatic mutations, comparing DNA and mRNA AI of driver mutations against a baseline of DNA and mRNA AI of somatic passenger mutations in the same genes, comparing across a large cohort of matched tumor genomes and transcriptomes. This approach does not require mRNA data from tumor-adjacent healthy tissues, and so enables a systematic study across all driver genes in 8, 809 tumors of 31 cancer types from the TCGA project. Recent studies of AI in somatic mutations focused on a single gene(8) or a single cancer type(8–14); we provide a comprehensive comparison of AI of 854, 785 somatic mutations across human cancers.

### Non-CNA AI contributes substantially to allelic imbalance in cancer transcriptomes

We hypothesized that mechanisms other than CNA may have a substantial impact on allelic mRNA expression imbalance of somatic variant alleles in cancer. To quantify the relative contributions of the well-understood CNA-AI and the less explored reg-AI towards the net mRNA-AI, we developed a framework of distinct beta-binomial models (see Methods).

First, an mRNA-AI model predicted RNA allele counts of a somatic mutation from RNA-Seq data, considering overall gene expression and sample tumor purity as confounders, to capture the overall imbalance of that mutation in the transcriptome, regardless of the cause. Second, to specifically assess CNA effects, we modelled DNA mutant allele counts from WES data using two models: one baseline model incorporated the sample purity, and the other the expected mutant copy number fraction (*CNF_mut_*, a quantity combining the variant allele frequency (VAF), purity and can) at the DNA level as a covariate (see Methods); this allowed us to determine if there is DNA-level AI beyond that expected from tumor purity, presumably attributable to CNA. Third, a reg-AI model was developed to assess allelic imbalance at the mRNA level beyond that attributable to genomic DNA dosage changes; this model predicted RNA allele counts, similarly to the mRNA-AI model, but incorporated the *CNF_mut_* as an additional covariate to correct for CNA effects.

For each somatic SNV, we determined significant AI by testing whether the observed variant allele frequency (*VAF_obs_*) significantly deviated from the model prediction (*VAF_pred_*) (see Methods). SNVs were classified into three categories: positive AI (pos AI) when *VAF_obs_* significantly exceeded *VAF_pred_* (upper-tail *p* < 0.05), indicating preferential expression of the *mutated* allele; negative AI (neg AI) when *VAF_obs_* was significantly below *VAF_pred_* (lower-tail *p* < 0.05), indicating preferential expression of the *wild-type* allele; and no significant AI (no AI) when neither condition was satisfied. This was done for each beta-binomial model separately to assess significant allelic imbalance by mRNA-AI, CNA-AI and reg-AI; the CNA-AI and reg-AI tests yielded mutually exclusive results by design (see Methods).

Of the 41, 549 somatic SNVs with significant mRNA-AI and additional significant reg-AI or CNA-AI in a pan-cancer analysis (5.6% of total SNVs), 46.6% exhibited reg-AI while 53.4% showed CNA-AI (Fig. 1B). This partitioning suggests that non-CNA mechanisms (reg-AI) might contribute for approximately half of mRNA-AI cases at these loci, with CNA-driven imbalance explaining the remainder. We note, however, that our reg-AI signal captures a composite of regulatory influences, alongside residual expression from contaminating normal cells (which we sought to mitigate in our models) and possible statistical artefacts. Similarly, we note that our CNA-AI estimate will also include effects of subclonal mutations, since they register as an imbalance at the DNA level similarly as the CNA do. Our analyses show this is responsible only for a minor part of significant CNA-AI calls in cancer genes (Figure S2) and moreover the subclonality can generate only negative CNA-AI, while our subsequent analyses focus on the positive AI (by either CNA-AI or reg-AI or overall mRNA-AI).

We observed differences between cancer types regarding the contribution of the reg-AI mechanisms on the overall mRNA-AI variability, ranging from 17.2% to 66.9% (Fig. 1B); however, in all cancer types the reg-AI effects on somatic mutations were considerable. Among the major cancer types where the CNA-AI was the predominant mechanism underlying the mRNA-AI were ovarian serous cystadenocarcinoma (OV, Fig. 1C), thyroid carcinoma (THCA [MIM: 188550]), brain lower grade glioma (LGG [MIM: 137800]), kidney renal papillary cell carcinoma (KIRP [MIM: 605074]) and pancreatic adenocarcinoma (PAAD [MIM: 260350]). On the other extreme, cancer types where the non-CNA mechanisms (reg-AI) were dominant contributors to mRNA-AI were the diffuse large B-cell lymphoma (DLBC), thymoma (THYM), uterine corpus endometrial carcinoma (UCEC, Fig. 1D), cervical squamous cell carcinoma (CESC), and skin cutaneous melanoma (SKCM [MIM: 155600]), reg-AI had the most influence on mRNA imbalance in expression of somatically mutated alleles. We note other major cancer types including stomach adenocarcinoma (STAD [MIM: 613659]), colon adenocarcinoma (COAD [MIM: 114500]), kidney renal clear cell carcinoma (KIRC [MIM: 144700]) and lung cancers (adenocarcinoma: LUAD; squamous cell carcinoma: LUSC [MIM: 211980]) also had a high contribution of reg-AI, suggesting widespread regulatory, possibly epigenetic mechanisms generating mRNA imbalances on somatic mutations.

Our comprehensive analysis of exome-wide somatic mutations in 31 cancer types reports higher estimates of non-CNA causes of mRNA-level AI than previous, focused studies. For instance, Correia *et al.*(8) attributed 84% of allele-specific expression in the *PIK3CA* gene in breast cancer to CNA effects. To validate these findings(8), we applied similar filters and data stratification (see Supplemental Material and Methods). This approach, focusing on breast invasive carcinoma (BRCA) and *PIK3CA* somatic variants was concurrent with previous analysis, where we estimated 16.7% (Wald 95% CI 0-37.8%) contribution from reg-AI and 83.3% (95% CI 62.2-100%) from CNA-AI, suggesting that even though reg-AI is fairly widespread, there are some genes where the relative role of CNA-AI is nonetheless more prominent. Another study of 1, 188 RNA-seq samples in 27 cancer types(15), quantified the relative contributions of various known factors to mRNA-level AI. The considered factors were germline eQTL (expression quantitative trait loci) variants, burden of proximal somatic SNVs, and the imprinting status of a genes (these known factors would, in our analysis, generate a part of the reg-AI signal); these explained 15.7% variance in their mRNA allelic imbalance, while the somatic CNA explained the rest(15). That this figure is smaller than our estimates of reg-AI suggest that other major effects, for instance resulting from epigenome remodeling in cancer(55–57), can generate mRNA-level AI on somatic mutations, and that they are more prevalent in cancer genomes than anticipated(8, 15).

For various downstream analyses in this study, we classified somatic SNVs in gene coding regions according to their effect. Specifically, we used the pathogenicity predictor *VARITY_ER*(44, 45) to reclassify low amino acid (aa) impact missense variants (score < 0.5, potentially non-pathogenic) and merge them together with the synonymous variants into an “effectively synonymous” (ES) mutation group, serving as a baseline. Additionally, we categorized genes based on their functional roles in cancer: oncogenes (OGs, n = 108), tumor suppressor genes (TSGs, n = 161), both TSG/OGs (n = 70) and passenger genes not found in Cancer Gene Census (CGC)(46) as a negative control (n = 15, 258), after filterings (see Methods). The *TP53* gene was treated independently, due to the large number of variants therein providing statistical power for individual analyses. We refer to “cognate genes” as those preferentially mutated in a specific cancer type, according to the MutPanning catalog(48). Therefore, the same driver genes are classified as cognate for some cancer types, while being noncognate for others.

### Mechanisms underlying regulatory allelic imbalance in cancer driver genes

The reg-AI signal in our data encompasses all types of AI variation at the mRNA level (mRNA-AI) that cannot be explained by significant DNA-level AI originating from CNA (CNA-AI). Therefore, mechanisms that give rise to reg-AI in cancer genes are of interest. We investigated the predicted impact of coding exonic mutations on splicing, which constitutes a post-transcriptional effect, and predicted mutation impact on promoter activity or chromatin status, which reflect a direct transcriptional effect (schematic in Figure 2A).

**Figure 2.**
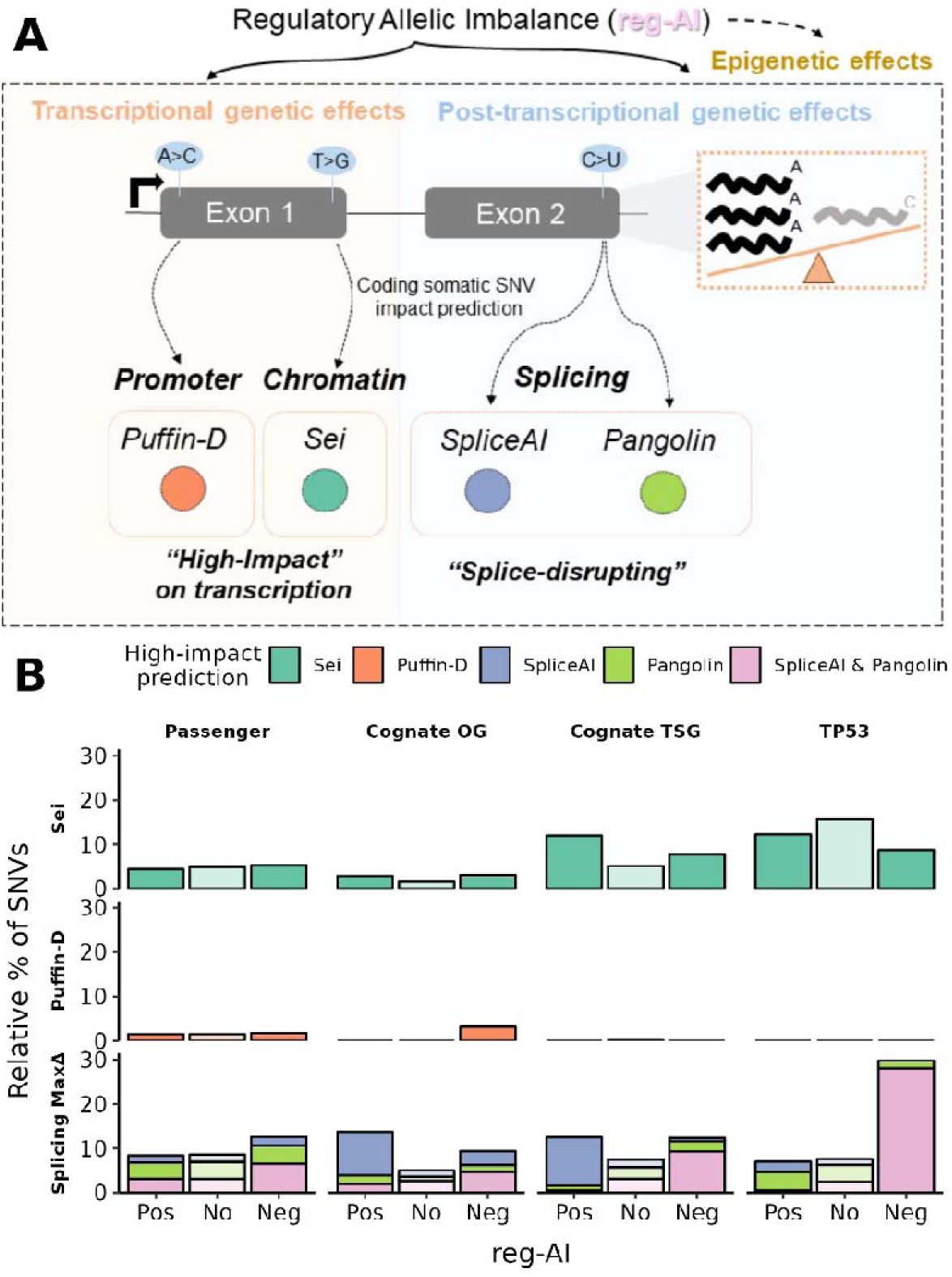
Mechanisms underlying regulatory allelic imbalance in cancer driver genes. **A,** Schematic illustrating the two primary mechanisms driving regulatory allelic imbalance in cancer driver genes: transcriptional and post-transcriptional genetic effects. We employed several computational tools to predict the functional impact of coding somatic SNVs on these processes. Transcriptional effects, which alter chromatin state or promoter activity, were assessed using *Sei*(65) and *Puffin-D*(62), respectively. Post-transcriptional effects on mRNA splicing were predicted using *SpliceAI*(58) and *Pangolin*(59). **B**, Proportions of somatic SNVs predicted to have a high impact through splicing disruption (Splicing MaxΔ), chromatin modification (*Sei*) or promoter activity (Δ*Puffin-D*), further stratified by gene classification and reg-AI status (see Supplemental Material and Methods). Splice disruption wa assessed using *SpliceAI* (*SP*_MaxΔ_ ≥ 0.15) or *Pangolin* (*PG*_MaxΔ_ ≥ 0.10) within ±50bp windows, with variants classified as splice-disrupting by either or both predictors. Chromatin effects were evaluated using *Sei* scores with the top 5% classified as high-impact, while promoter effects were assessed using Δ*Puffin-D* with high-impact variants defined as those exceeding the first quartile of the positive control variants (*TERT* somatic mutations known to affect promoter activity(63, 64)). All these proportions are shown for distinct gene categories: passenger genes, cognate oncogenes (OGs), cognate tumor suppressor genes (TSGs) and *TP53* analyzed independently. We refer to “cognate genes” as those preferentially mutated in a specific cancer type, according to the MutPanning catalog(48). Therefore, the same driver genes are classified as cognate for some cancer types, while being noncognate for others. Within each gene category, SNVs are further grouped by reg-AI significance: positive reg-AI (indicating preferential expression of the mutated allele), no reg-AI (no significant AI at the mRNA level attributable to regulatory effects), and negative reg-AI (preferential expression of the wild-type allele).

First, we evaluated every coding variant with two deep-learning predictors—*SpliceAI*(58) and *Pangolin*(59)—classifying a mutation as “splice-disrupting” when its maximal Δ-score was ≥ 0.15 (*SP*_MaxΔ_) or ≥ 0.10 (*PG*_MaxΔ_) respectively, within a ±50-bp window (see Supplemental Material and Methods). For instance, among cognate TSGs, 4.9% of mutations with no significant reg-AI were splice-disrupting by *SpliceAI* (Fig. 2B), compared to 10.9% of those with significant reg-AI (OR = 2.4, *p* = 4.1e-5, Fisher exact test; positive reg-AI: 11.5%; negative reg-AI: 10.1%). For cognate OGs we observed a similar pattern (4.1% no AI vs 10.2% reg-AI, OR = 2.7, *p* = 1.3e-3; positive reg-AI: 11.8%; negative reg-AI: 7.8%). Therefore, approximately 6% of significant reg-AI of coding somatic mutations in cancer genes can be explained by these mutations causing splicing disruption. Subsequent enrichment analyses confirmed signatures of selection for splice-mediated loss-of-function effects of coding mutations in TSGs (negative reg-AI) and gain-of-function in OGs (positive reg-AI), depending on the specific sets of SNVs with distinct functional impact: ES, high or low aa-impact missense, synonymous-only (detailed analysis provided in Text S1 and Figure S3). This substantiates previous reports of splice-altering coding driver mutations in OG and TSG(60, 61) by additional evidence from mRNA allelic imbalances. As a prominent example, we examined 3 known recurrent splice-altering exonic variants in *TP53*(60), 2 synonymous and 1 missense, which commonly resulted in negative reg-AI but not CNA-AI (Text S1, Figure S3A). As a side note, we observed an association of mRNA-level AI with splice-altering missense mutations as well as synonymous ones, suggesting that splicing effects could be an overlooked mechanism for missense cancer driver mutations (Text S1). Overall, splicing effects of somatic mutations do cause reg-AI, and reg-AI can be used to prioritize the splice-altering coding mutations in cancer, however the proportion of cancer reg-AI explained by splicing effects of coding point mutations is modest.

Next, we considered whether the coding mutations could underlie a direct transcriptional modulation, thereby being a cause of reg-AI. Using the *Puffin-D* deep-learning predictor(62) to assess effects on promoter activity (see Supplemental Material and Methods), we found that only 0.6% of coding variants in cancer genes (all TSGs and OGs) had Δ*Puffin-D* scores exceeding the first quartile of the positive control variants (*TERT* somatic mutations known to affect promoter activity(63, 64)) (Fig. 2B and Figure S4). Therefore, when considered in aggregate, the effects of coding SNVs on reg-AI via this mechanism cannot be substantial. To assess impacts on local chromatin, we used the *Sei* deep-learning predictor of local chromatin state(65), and classified the top 5% of SNVs with highest scores in affecting the *Sei* active transcription states (TN1-TN4) as potentially impactful (positive controls are lacking). This analysis revealed an enrichment among only cognate TSGs, where 12% of mutations exhibiting positive reg-AI had high *Sei* scores compared to 5% with no reg-AI (OR = 2.50, *p* = 4.6e-4, Fig. 2B), suggesting an interesting example of apparently gain-of-function mutations in TSG that also increase mRNA-level of the *mutant* allele. The other considered cancer gene categories did not have significant effects by *Sei* score, therefore also this analysis suggests that direct effects of somatic coding SNV on transcriptional output are uncommon. Detailed analysis and discussion of *Puffin-D* and *Sei* predictions linked with reg-AI is provided in Text S2.

While direct transcriptional effects of coding mutations proved rare, we considered whether reg-AI might instead result from additional non-coding somatic mutations affecting regulatory regions, analogous to how germline eQTL variants cause allele-specific expression by altering transcription factor binding sites. To test this possibility, we analyzed TCGA tumors with WGS data available (n=777) for co-occurring promoter-proximal mutations (±1.5kb from TSS) in the same tumor samples as cancer driver genes bearing exonic variants exhibiting significant reg-AI (Figure S5). Among these coding mutations, 55% of those with positive reg-AI had co-occurring promoter-proximal non-coding mutations, compared to 44% of mutations without reg-AI, suggesting that approximately 11% of positive reg-AI events in cancer driver genes result from nearby regulatory non-coding mutations (OR = 1.54, *p* = 6.8e-2). In contrast, passenger genes did not show this pattern, with actually fewer promoter mutations in positive reg-AI cases, suggesting positive selection on promoter mutations in drivers. Detailed analysis is also provided in Text S3. A recent study of 1, 188 cancer RNA-Seq samples(15) did also find an association of promoter mutations on mRNA allelic imbalance, with a modest effect size, below splice-region mutations.

Collectively, our analyses above indicate that splice-disrupting exonic variants and also TSS-proximal non-coding variants can explain some cases of reg-AI for somatic mutations in cancer genes, while direct transcription-altering mechanisms of mutations play a minor role.

### Driver tumor suppressor genes show preferential expression of mutated alleles in high aa-impact missense variants

Above, we established that splice-altering coding mutations and also promoter-proximal non-coding mutations are under positive selection and can explain a certain component of reg-AI in cancer driver genes. The majority of reg-AI instances are not explained by these mutational causes (nor, by definition of reg-AI, explained by CNA) suggesting they are caused by allele-specific epigenetic changes. Next, we turn from studying the causes of the reg-AI, to ask whether selection acts on reg-AI and overall mRNA-AI more broadly, regardless of the cause of the reg-AI. We assessed across all somatic coding variants in cancer genes, whether reg-AI can effectively generate second-hit events similarly as CNAs commonly do(4, 5), by using the synonymous plus low aa-impact missense mutations (both grouped as effectively synonymous (ES)) as a neutral baseline.

To evaluate whether AI upon mutations is selected in cancer genes we assessed whether the proportion of high aa-impact missense variants (compared to effectively synonymous) was significantly higher for mutations associated with significant AI (either positive or negative, in separate enrichment tests), for each gene category (Fig. 3A, see Methods). For nonsense variants, we analyzed their proportions in comparison to the other two SNV types.

**Figure 3.**
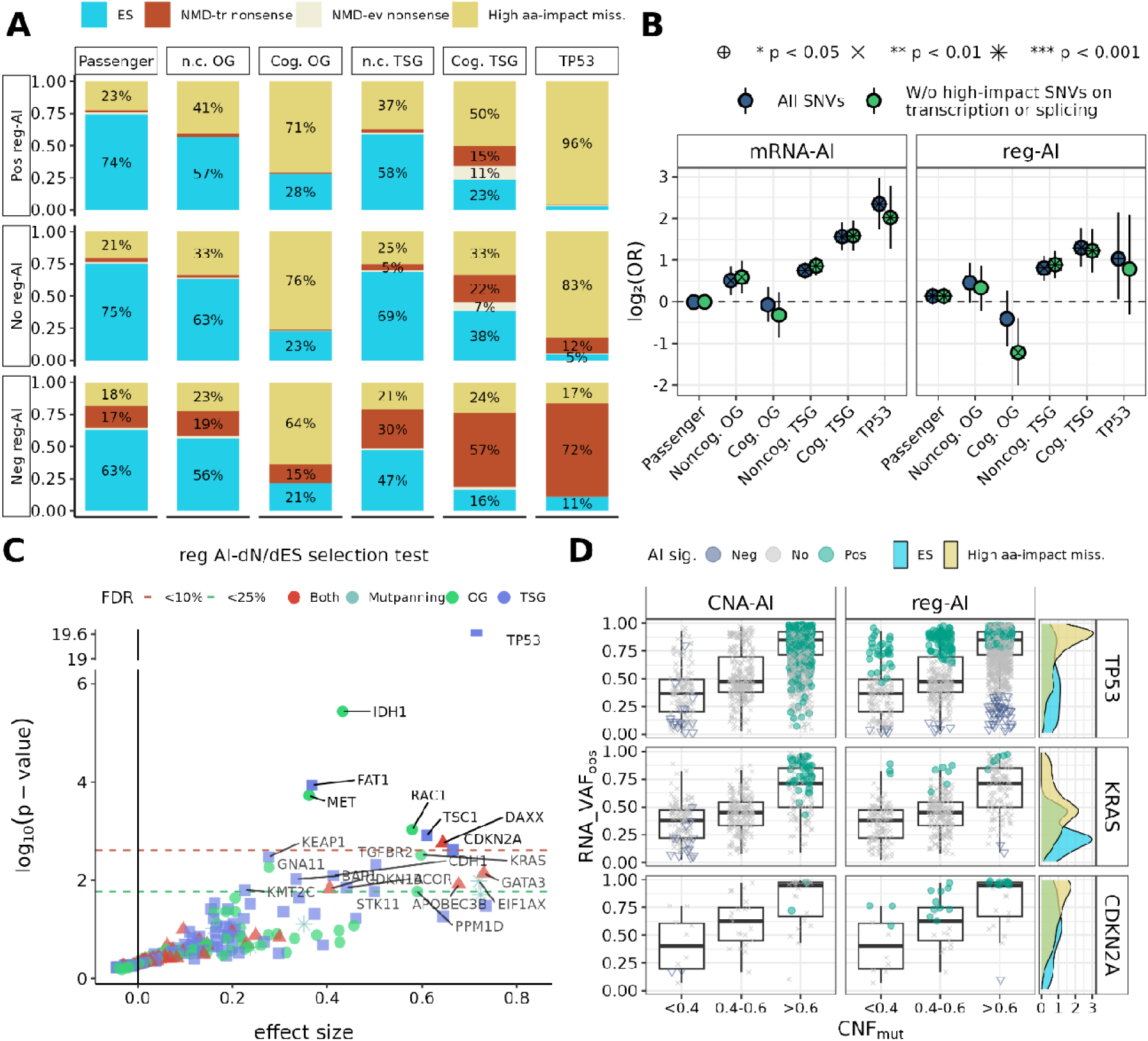
Allelic imbalance is subject to selection in cancer transcriptomes, potentially contributing to tumorigenesis. **A**, Relative frequencies of somatic coding SNVs, categorized by their predicted functional impact: effectively synonymous (ES), high amino-acid (aa)-impact missense and nonsense. Nonsense mutation were further subdivided into those predicted to trigger nonsense-mediated mRNA decay (NMD-tr) and those predicted to evade (NMD-ev) based on known genomic rules(67) (see Supplemental Material and Methods). Proportions were further stratified by gene categories: passenger genes, OGs, TSGs and *TP53* (analyzed independently). OGs and TSGs were additionally split into cognate (Cog.) and noncognate (n.c.). “Cognate” means SNVs matched to the cancer type where they are known to be a driver based on MutPanning catalog(48). Within each gene and functional impact category, proportions are grouped based on their reg-AI significance: no reg-AI (indicating no significant allelic imbalance at the mRNA level attributable to regulatory effects), positive reg-AI (indicating preferential expression of the *mutated* allele) and negative reg-AI (indicating preferential expression of the *wild-type* allele). A higher proportion of positive reg-AI among missense mutations is indicative of selection pressure favouring increased mRNA expression of these *mutant* alleles. **B,** Enrichment analysis of high aa-impact missense mutations exhibiting positive AI (mRNA-AI left panel, reg-AI right panel) stratified by gene category. ORs and 95% CIs are presented in log_2_ scale. Within each gene category, enrichments are estimated for (1) all somatic SNVs and (2) excluding mutations that are also predicted to be splice-disrupting or impact transcription regulation based on stringent thresholds (*SpliceAI SP*_MaxΔ_ ≥ 0.10; *Pangolin PC*_MaxΔ_ ≥ 0.10; Δ*Puffin-D* scores above the 25th percentile of positive controls; *Sei* scores from the top 10th percentile). Statistical significance was assessed using Fisher’s Exact Test, with *p*-values indicated as: \**p* < 0.05, \*\**p* < 0.01, \*\*\**p* < 0.001. **C**, Volcano plot illustrating the results of our gene-level reg AI-dN/dES test for 356 total genes: OGs, TSGs, genes classified as both TSG/OGs, and MutPanning-only genes. The test adapts our pan-cancer beta-binomial framework, originally for detecting significant AI at the single SNV-level, to assess gene-level reg-AI (see Methods). The X-axis displays the effect size of the regression coefficient for the *SNV_type_* covariate (high aa-impact missense versus ES mutations) from the model. The Y-axis represents the raw upper-tail *p*-values from this regression, presented on a -*log*_10_ scale. Genes exhibiting a statistically significant effect (FDR < 10%) or candidates (FDR < 25%) are highlighted, with its corresponding thresholds as horizontal dashed lines. A positive and statistically significant effect size for a gene indicates that its high aa-impact missense mutations are associated with a greater enrichment of the mutant allele’s mRNA expression compared to the neutral baseline established by its ES mutations. This signature suggests that the observed AI itself is under positive selection in these identified driver genes, potentially contributing to their oncogenic role. **D**, *RNA_VAF_obs_* distribution of SNVs plotted against estimated mutant copy number fraction (*CNF_mut_*, see Methods) for three positively selected genes (*TP53*, *KRAS*, *CDKN2A*), stratified by AI significance status (positive, no significance, or negative) for CNA-AI, mRNA-AI, and reg-AI. *CNF_mut_* is a proportion value stratified in three bins (<40%, 40-60%, >60%). Individual SNVs may exhibit significance for mRNA-AI alongside either CNA-AI or reg-AI, though concurrent significance for both CNA-AI and reg-AI is uncommon. As expected, SNVs significant for CNA-AI typically correlate with higher *CNF_mut_*, while those significant for reg-AI show the independence from copy number alterations. *TP53* displays numerous SNVs significant for both reg-AI and CNA-AI, demonstrating dual mechanistic contributions to overall mRNA-AI; *KRAS* shows predominantly CNA-AI significant SNVs with fewer reg-AI events; *CDKN2A* exhibits primarily reg-AI significant SNVs. Across all three genes, high aa-impact missense mutations consistently demonstrate higher *RNA_VAF_obs_* than ES mutations, supporting their identification as positively selected in the AI-dN/dES analysis (panel C).

First and as expected, nonsense mutations across all gene categories were enriched for variants associated with negative AI, (mRNA-AI: OR = 3.59, CI 95% [3.47-3.72]; reg-AI: OR = 6.12, CI 95% [5.93-6.3], both *p* < 2e-16, Fisher’s exact test, Fig. 3A). This is explained by NMD, a surveillance pathway that degrades mRNAs containing premature termination codons, a well-known cause of allele-specific expression(26, 66–68). The enrichment was primarily observed for nonsense variants predicted to trigger NMD (Fig. 3A) based on established genomic rules(67) (see Methods), particularly high for *TP53* (reg-AI: OR = 19.3, CI 95% [13.68-27.60], *p* < 2.2e-16), compared to other cognate TSGs (reg-AI: OR = 4.64, CI 95% [3.64-5.94], *p* < 2.2e-1; *TP53* excluded) or passenger genes (reg-AI: OR = 7.6, CI 95% [7.29-7.82], *p* < 2.2e-1). Nonsense mutations predicted to escape NMD showed the opposite trend (Fig. 3A): in cognate TSGs, their prevalence rose from 2% among SNVs with negative reg-AI, to 7% with no reg-AI, and to 11% with positive reg-AI (OR = 1.5, *p* = 5e-2). This stepwise enrichment of NMD-evading nonsense mutations highlights truncated proteins that may exert dominant-negative or gain-of-function effects, and therefore have the mutant allele preferentially expressed.

In subsequent comparisons we contrast the impactful missense versus a baseline of ES mutations (“dN/dES” test, see Methods), while excluding nonsense mutations when calculating enrichments (odds ratios, ORs below) to avoid confounding by NMD on AI. Testing our main hypothesis, we considered cognate TSGs, where the prevalence of high aa-impact missense mutations rose from 33% in tumors with no reg-AI to 50% in tumors exhibiting positive reg-AI (Fig. 3A) (OR = 2.45, CI [1.78-3.41], *p* = 6.3e-9, Fig. 3B) and from 31% with no mRNA-AI to 52% in examples with significant pos mRNA-AI (Figure S6A) (OR = 2.95, CI [2.35-3.73], *p* = 8.7e-23, Fisher test, Fig. 3B). The same pattern held for *TP53*, which had sufficient number of mutations to be assessed individually: relative prevalence of impactful mutations increased from 61% to 95% for mRNA-AI (OR = 5.08, CI [3.35-7.85], *p* = 1.2e-16) and from 83% to 96% for reg-AI (OR = 2.04, CI [1.05-4.45], *p* = 3.3e-2). Interestingly, this enrichment was also observed for (supposedly) non-cognate OGs and TSGs, suggesting that reg-AI could identify additional driver genes relevant to a certain cancer type.

Cognate OG, considered collectively, did not show association of mRNA-level AI with impact of missense mutations (Fig. 3B), which does not preclude that individual genes could show the AI association. As a control, passenger genes showed no significant differences in enrichment between positive mRNA-AI and no mRNA-AI for the impactful missense mutations (*p* = 0.67), albeit there was a slight background enrichment for reg-AI (OR = 1.10, CI [1.05-1.14]) which represents a modest bias in our methodology and/or presence of some weaker driver genes in the passenger gene group.

Next, we aimed to rule out that these missense variants, selected via occurrence positive reg-AI, directly caused the AI rather than acting, as we hypothesize, in a two-hit mechanism involving the coding driver mutation and another, presumably epigenetic event causing AI. After applying stringent filters to exclude splice-disrupting variants (*PG*_MaxΔ_ ≥ 0.1 or ≥*SP*_MaxΔ_ ≥ 0.1) and mutations impacting transcriptional activity (Δ*Puffin-D* scores above the 25th percentile of positive controls and *Sei* scores from the top 10th percentile), the enrichment signals persisted (Fig. 3B), only modestly attenuated for cognate TSGs (mRNA-AI OR = 3; reg-AI OR = 2.33) and *TP53* (mRNA-AI OR = 4.05; reg-AI OR = 1.72). These findings demonstrate that regulatory AI selection in cancer driver genes mainly operates through mechanisms beyond direct effects of driver coding mutations on transcriptional mechanisms (promoter output, chromatin state) or post-transcriptional processes (splicing).

In summary, missense variants in cancer driver genes showing preferential expression of the mutant allele are subject to stronger positive selection than driver gene variants not showing AI of the mutant allele, highlighting the key role of mRNA-AI as second-hit events in tumor evolution. Our test of selection, when applied to reg-AI, further suggests that the non-CNA causes of mRNA-level allele expression imbalance can have a strong effect on selection on driver mutations and should be considered alongside CNAs in the ability to generate driver mutations. In particular, AI correlated with the *VARITY* functional impact scores for missense mutations in cognate TSGs (including *TP53*), at the reg-AI level (Pearson’s R = 0.23) and at the mRNA-AI level (R = 0.4; Figure S6B).

#### Non-CNA mRNA AI in individual driver genes is subject to positive selection

To further identify individual cancer driver genes subject to selection on mRNA allelic imbalance, we developed AI-dN/dES, a dN/dS-type test (see Methods). AI-dN/dES is based on our previous pan-cancer beta-binomial framework for single-SNV significant AI detection, now applied at the gene level, separately for mRNA-AI and reg-AI. The model incorporates an *SNV_type_* covariate, and its regression coefficient measures the gene-specific differential AI between missense and ES mutations: a positive, statistically significant effect size (by upper tail *p*-value) indicates that high aa-impact missense mutations drive greater enrichment of the *mutant* allele in mRNA than the neutral baseline of ES variants. This signature provides direct evidence that AI itself is under positive selection in the identified driver genes. We performed AI-dN/dES tests for mRNA-AI and separately for the reg-AI. Firstly, a genome-wide search for reg-AI dN/dES significance (n = 10, 817) resulted in a 2.5-fold (at FDR 10% gene-wise significance) and 2.9-fold (at FDR 25%) enrichment of known cancer driver genes. When considering the overall mRNA-AI, which additionally captures the CNA signal, the enrichments were expectedly increased: 10.4-fold (at FDR 10%) and 4.9-fold (at FDR 25%). Among the putative passenger genes that we identified at FDR<10% (listed in Table S2), this mRNA-level analysis in mutation imbalance suggests a possible driver role (26 genes for mRNA-AI; 39 genes for reg-AI).

Next, we focused on our set of known cancer driver genes from CGC to reduce the multiple-testing burden, additionally including 16 passenger genes identified by MutPanning(48) but not found in CGC, leading to a total of 356 genes for testing after filterings (see Methods).

At FDR < 10%, we identified significant positive selection on AI favoring the mutant allele in 18 genes for mRNA-AI (Figure S6C) and 8 genes for reg-AI (Fig. 3C). Relaxing the threshold to FDR < 25 % expanded these sets to 35 and 22 genes for overall mRNA-AI and specifically the reg-AI, respectively (listed in Table S3). Q-Q plots indicated that the *p*-values are broadly well calibrated (Figure S6D). To assess robustness, we performed a sensitivity analysis excluding subclonal variants, which can potentially confound selection estimates due to incomplete representation across tumor cell populations; they also register as negative CNA-AI by our methodology although mRNA-AI estimates are largely unaffected. This analysis yielded qualitatively consistent results at FDR<25% (Figure S6E): 39 genes for mRNA-AI (30 genes, 86% overlapping with the original analysis) and 19 genes for reg-AI (17 genes, 77% overlapping). The modest reduction in significant genes reflects decreased statistical power from the smaller variant set rather than systematic bias.

Tumor-suppressor genes were more common hits than oncogenes (mRNA-AI: 12 TSGs vs 4 OGs; reg-AI: 4 TSGs vs 3 OGs), as anticipated from our previous enrichment analysis (Fig. 3A and Figure S6A). Seven of the 8 reg-AI hits, *TP53*, *IDH1*, *FAT1*, *TSC1*, *CDKN2A*, *MET* and *RAC1*, also emerged from the mRNA-AI analysis as expected as mRNA-AI also encompasses reg-AI mechanisms; *DAXX* was unique to reg-AI. *TP53* exhibited the strongest signal: high aa-impact missense mutations increased the log-odds of mutant allele expression by 0.72 in reg-AI (FDR = 1e-17) and by 0.93 in mRNA-AI (FDR = 1e-23), corresponding to ∼2-2.5-fold higher odds of missense relative to ES variants (Fig. 3C). Across the remaining significant genes, effect sizes ranged from OR = 1.43–1.93 in reg-AI and from OR = 1.43–3.52 in mRNA-AI, underscoring positive selection on AI in diverse cancer driver genes. By contrast to the above, 11 genes (*KEAP1*, *BAP1*, *STK11*, *EIF1AX*, *KRAS*, *TGFBR2*, *CUL3*, *NF1*, *PPP2R1A*, *CIC* and *SMAD3*) were significant only in the mRNA-AI analysis but not in reg-AI at FDR < 10%. Notably, of those, *KEAP1*, *BAP1*, *STK11*, *EIF1AX*, *KRAS* and *TGFBR2* are reg-AI significant at FDR < 25% (Fig. 3C), underscoring that the two mechanisms for generating mRNA allelic imbalance, the CNA-dependent and the CNA-independent one, affect a largely overlapping set of genes.

However, genes can be affected by the two AI mechanisms with different frequencies. For example, in *KRAS*, high aa-impact missense variants exhibited higher *RNA_wAF_obs_* than ES variants (Fig. 3D), with a substantially larger fraction of SNVs reaching significance under the CNA-AI compared to the reg-AI criteria (12.5% vs 4.1%, respectively). Conversely, in *CDKN2A*, positive selection on putative driver SNV was driven predominantly by reg-AI-specific variants (7.14% vs 46.42%), while in *TP53* the contributions were more balanced between the two mechanisms (29% vs 20%) (Fig. 3D).

Tumor suppressor genes are known to frequently undergo LOH through copy-number deletion of the *wild-type* allele, with the somatic mutation serving as the second hit. To validate our analytical framework under LOH events, we examined our 12 selected TSGs via mRNA-AI and compared AI patterns between LOH (*wild-type* allele copy number, *ACN_WT_* = 0) and non-LOH cases (*ACN_WT_* >= 1, see Methods). As expected, *RNA_VAF_obs_* was substantially higher in LOH cases compared to no LOH cases (Figure S7), with a bigger increase for high aa-impact missense variants than ES variants, consistent with the two-hit model via LOH. Among LOH cases, 35% registered as significant for positive CNA-AI in our tests, whereas only 18% exhibited positive reg-AI (Figure S7). Thus, most mRNA allelic imbalance in LOH cases is adequately explained by copy number changes, rather than regulatory mechanisms, expectedly. Conversely, non-LOH cases of these 12 TSGs showed minimal positive CNA-AI (0.92%) but higher rates of positive reg-AI (6.56%), suggesting that regulatory mechanisms become more prominent in TSGs, in relative terms, when the *wild-type* allele remains present. In other words, for a cancer gene, at the population level, the CNA-AI and reg-AI mechanisms affect overlapping sets of genes, however at the level of an individual patient there is a mutual exclusivity between the mechanisms of gaining AI.

Similarly as for TSGs above, we turned to examine oncogenes such as *KRAS*, which can, in many cases, be activated through copy-number amplification of the mutated allele(4). We examined 8 OGs showing significant mRNA-AI (relaxed FDR<25%) and compared AI patterns between *CN_neut_* (copy-number neutral) (*ACN_MUT_* = 1 and *ACN_WT_* = 1) and *CN_gain_* (copy-number gained) (*ACN_MUT_* >1, see Methods). *RNA_VAF_obs_* was modestly higher in *CN_gain_* versus *CN_neut_* cases, and the increase was larger in high aa-impact missense variants than in ES variants (Figure S8). Among *CN_gain_* cases, 14% of SNVs showed significant positive CNA-AI, and only 2% exhibited positive reg-AI (Figure S8), supporting a mutually exclusive occurrence in OGs like in TSGs. Conversely, the many *CN_neut_* cases showed minimal positive CNA-AI (0.23%) but 3.6-fold higher albeit still modestly common positive reg-AI (0.82%), suggesting an occasional role for regulatory AI mechanisms in copy-neutral contexts of some OGs.

Overall, allelic expression imbalance favoring the allele harboring a somatic driver mutation confers a selective advantage in various cancer driver genes. Non-CNA mechanisms are an important, and in some genes the dominant mechanism in generating the mRNA-level AIs that are under positive selection in somatic evolution.

### Positive selection on AI adversely impacts cancer prognosis

Given the signatures of positive selection on AI of driver somatic mutations, we hypothesized that this would be reflected in cancer aggressiveness. To test this, we performed overall survival analyses, in which we compared patients with tumors exhibiting AI on driver mutations in genes where we identified significant selected AI, against patients with tumors that bore mutations in the same drivers but with no detectable AI.

For this analysis, we selected only samples bearing at least one high aa-impact missense mutation within any of the genes previously identified as having significant positive selection for mRNA-AI (34 genes at FDR < 25%; permissive threshold to increase sample and variant retention and boost statistical power); ES variants were excluded. *TP53* was considered separately here, to avoid bias from its exceptionally high mutation frequency.

To quantify the extend of AI for each tumor, we first calculated the Δ*RNA_VAF* as the difference between the *RNA_VAF_obs_* and the *RNA_VAF_pred_* by our beta-binomial models for each qualifying SNV (see Methods). The mean of these Δ*RNA_VAF* across all considered SNVs within a given tumor was then used as a continuous AI score for that sample. Tumor samples were subsequently classified into four categories based on this continuous AI score: neutral/reference ([-0.1, 0.1]); and positive AI across three quantile-based bins (for mRNA-AI: (0.1, 21], (0.21, 0.37] and (0.37-0.7]).

In the Kaplan-Meier (KM) curves for our pan-cancer analysis, patients within the positive mRNA-AI affected groups had a significantly poorer prognosis (*p* = 2.95e-6, log-rank test), where 35.4% of the affected patients have died during the follow-up period, while only 25.2% of the non-affected patients did (Fig. 4A). Similar results were observed for reg-AI (Figure S9A, *p* = 9.45e-6) using its corresponding set of positively selected AI genes at FDR < 25% (n = 21, excluding *TP53*). Considering the two AI analyses, the median overall survival for the AI affected group was 3.68 – 3.78 years, compared to 7.96 – 8.18 years for the AI non-affected group.

**Figure 4.**
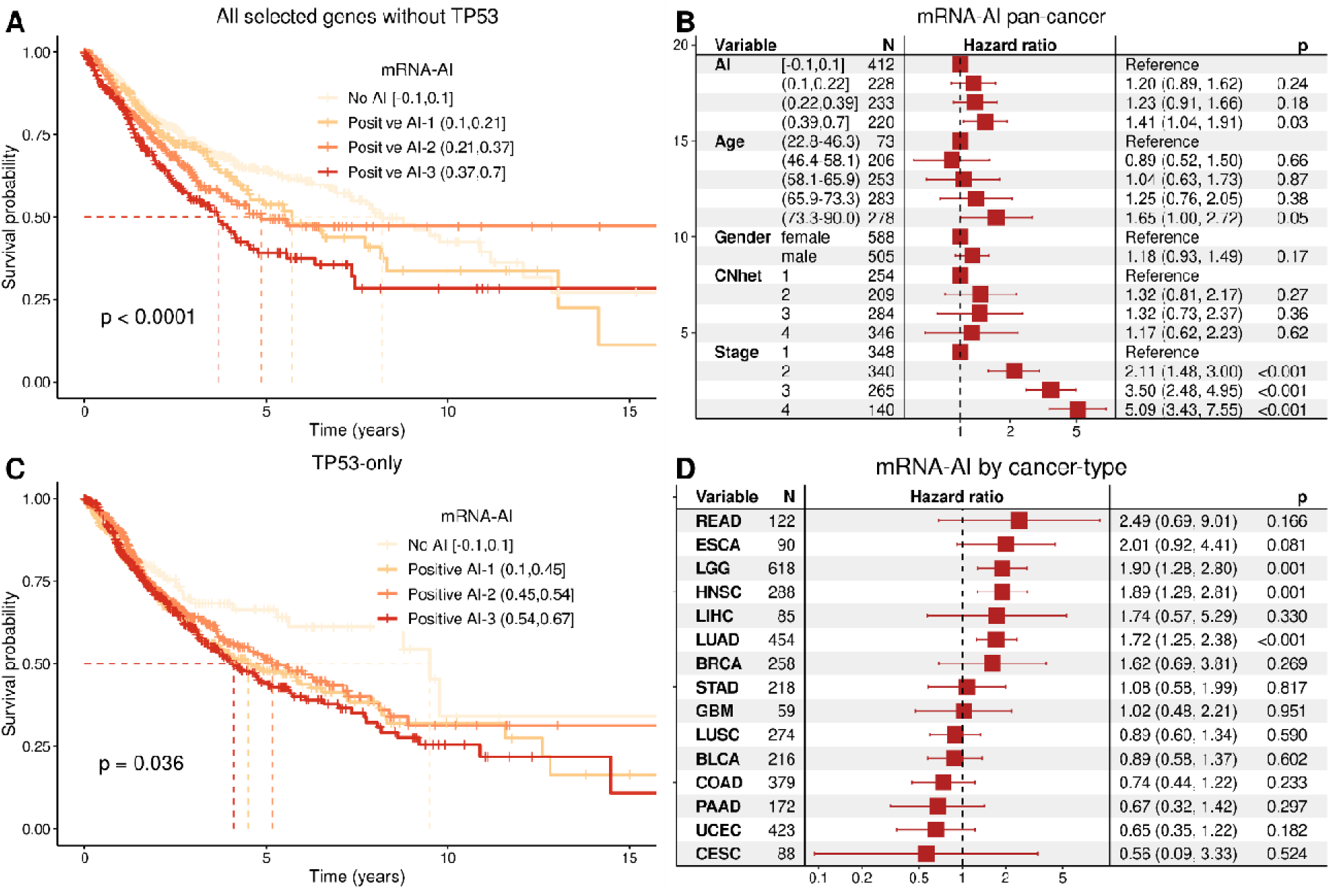
Positive selection on allelic imbalance has a detrimental effect on overall cancer prognosis. **A**, Kaplan-Meier (KM) overall survival curves for pan-cancer patients harbouring at least one high aa-impact missense mutation in 35 genes under positive mRNA-AI selection (FDR < 25%; *TP53* excluded; ES mutations excluded). Patients are stratified by per-sample mRNA-AI score into three “positive” quantile bins groups ((0.1, 0.21], (0.21, 0.37], or (0.37, 0.7]) versus “neutral/reference” ([–0.1, 0.1]). This mRNA-AI score is calculated based on the, calculated as the difference between the and the by our beta-binomial model (see Methods) for each qualifying SNV. The mean of these across all considered SNVs within a given tumor was then used as a continuous mRNA-AI score for that sample. The indicated *p*-value is derived from a log-rank test. **B,** Co proportional hazards survival analysis, showing the estimated hazard ratios (HR) for each variable, along with the 95% CIs and *p*-values. The results demonstrate a significant positive association between increasing mRNA-AI intensity and higher HRs. The model incorporated the following covariates: patient age (discretized into quintiles), sex, tumor stage, cancer type, intratumor genetic heterogeneity (estimated as copy-number heterogeneity, CNH), tumor sample purity, and total sample CNA burden. Some of these covariates are excluded from the plot for better visualization. **C**, KM curves for pan-cancer overall survival, analogous to the analysis presented in panel A for mRNA-AI, but exclusively considering patients with high aa-impact missense mutations in *TP53*. **D**, Cox proportional hazards survival analysis, analogous to panel B, performed separately for individual cancer types. Due to varying sample sizes across cancer types, a modified approach was employed using binary AI classification based on SNV-level significance from our beta-binomial models. Tumors were stratified as AI “affected” (harboring at least one high aa-impact SNV with significant positive AI) versus AI “not affected” (harboring SNVs without significant positive AI; cases with negative AI excluded). This analysis included *TP53* mutations to enhance statistical power and was applied to 15 cancer types meeting predefined filtering criteria (see Methods). Cancer types with fewer than 50 samples were excluded from analysis.

Because this analysis does not consider cancer type or other covariates, we next conducted a Cox proportional hazards survival analysis (see Methods), incorporating cancer type and additional covariates known to be associated with cancer survival (Fig. 4B). Covariates used in the model included patient age (discretized into quintiles), sex, and tumor stage. Additionally, the model also accounted for other relevant variables known to predict survival, in particular the cancer type, intratumor genetic heterogeneity (estimated as copy-number heterogeneity, CNH score)(54), tumor sample purity and total CNA burden. After controlling for these covariates, we still observed that patients categorized in the positive mRNA-AI ranges faced a progressively higher risk of death, compared to mRNA-AI non-affected patients, with hazard ratios (HR) ranging from 1.20 to 1.41 (95% CI 0.89-1.04, 1.62-1.91; *p* = 0.24 - 3e-2), where higher HRs were positively associated with higher degree of AI (Fig. 4B).

The effect size of positive mRNA-AI signal is quite substantial in comparison to known survival covariates: the HR of CNH was of 1.32 for the highest quartile compared to the lowest (*p* = 0.27). Patients within the 4th and 5th age quintiles, ranging from 66-73 and 73-90 years old, exhibit hazard ratios comparable to those seen with our mRNA-AI variable, ranging from 1.25 to 1.65 (*p* = 0.38 to 5e-2). This implies that the prognostic impact of somatic mutation AI in tumor transcriptomes on cancer patient survival is comparable to the effect of being 27-50 years older (compared to our youngest reference age group, which was 23-46 years old). We observed similar results specifically for the reg-AI signal across the three positive AI categories (HR = 1.17-1.52; *p* = 0.42 – 2.2e-2, Figure S9B), suggesting that the non-CNA mechanisms driving AI, also contribute to cancer aggressiveness.

#### AI in individual genes and cancer types linked with cancer aggressiveness

Additionally, we asked if the association of AI with survival held for individual driver genes (from our positively selected gene sets) and cancer types (of those where a high number of mutations/samples was available to facilitate testing). For example, considering only patients harboring at least one *TP53* high aa-impact missense variant, those within positive mRNA-AI categories in *TP53* had a poorer prognosis than those who were not affected by AI: (Cox HR = 1.44-1.62, *p* = 9.7e-2 – 2.2e-2; KM curve in Fig. 4C). It was non-significant for *TP53* reg-AI (HR = 1.08-1.21 with *p* = 0.21-0.63, KM curve in Figure S9C). Analyses of other individual drivers were generally underpowered or yielded non-significant results.

Next, we asked in which cancer types the AI-survival associations were more prominent. In this case, the number of mutations and available sample sizes varied per cancer type, and we adapted our previous approach. For this, we employed a binary AI classification based on SNV-level significance derived from our beta-binomial models. Tumors were categorized as being AI “affected” (harboring SNVs with significant positive AI) versus AI “not affected’ (harboring SNVs without significant positive AI, and excluding cases with negative AI). This strategy, which included *TP53* data to enhance statistical power, was applied to 15 cancer types that met our filtering criteria (see Methods). Positive mRNA-AI was significantly prognostic (FDR < 5%) in head and neck squamous cell carcinoma (HNSC; mRNA-AI HR = 1.89, FDR = 8.4e-3; reg-AI HR = 0.79, ns), lung adenocarcinoma (LUAD; mRNA-AI HR = 1.72, FDR = 8.4e-3; reg-AI HR = 1.23, ns), and low-grade glioma (LGG; mRNA-AI HR = 1.90; reg-AI HR = 0.69, ns) (Fig. 4D).

### Suggestive evidence for an epigenetic cause in non-CNA allelic imbalance

Our analyses above suggest that reg-AI of coding somatic mutations, in most instances, results from mechanisms beyond splicing-altering or direct transcriptional/chromatin effects of the coding mutation, or of co-occurring noncoding mutations in the gene promoter. Epigenomic inactivation, for example, is a known second-hit event for germline cancer risk variants in colon and breast cancers(3), therefore, in principle, somatic mutations might be affected by similar epigenetic two-hit mechanisms. To support that epigenetic causes underlies this CNA-independent allelic expression imbalance phenomenon (reg-AI), we sought signatures of the differential activity of DNA regulatory elements between alleles.

To examine them, we used matched coding WES and ATAC-Seq data from 381 TCGA patients(35) and we performed variant calling on the ATAC-Seq reads to estimate allele-specific chromatin accessibility (ASCA) for *mutant* alleles. Then, we applied the same beta-binomial modelling approach used for RNA-Seq data, allowing us to test whether allelic imbalances in the chromatin accessibility would be able to explain allelic imbalances observed at the mRNA level. Considering variants with stringent coverage threshold (> 20 total reads) in both RNA-Seq and ATAC-Seq, we analyzed 782 somatic variants that overlapped with ATAC-Seq peaks of sufficient intensity, meaning they were located in or near to active CREs within the gene (e.g. intragenic enhancers).

We examined the correlation between ASCA and reg-AI by comparing the residuals from our copy number-corrected models (Δ*ATAC_VAF* and Δ*ATAC_VAF*, see Supplemental Material and Methods) for both datasets. ASCA indeed significantly predicted reg-AI for high aa-impact missense variants (Pearson’s R = 0.45; *p* = 1.9e-89), and effectively synonymous variants (R *=* 0.47*; p =* 2.2e-16) (Figure S10A). Expectedly, this correlation did not extend to nonsense variants (R = 0.2; *p* = 0.33): due to the degradation of transcripts containing PTCs through NMD, transcript abundance is uncoupled from the effects of chromatin accessibility on transcription. This pattern remained consistent across different coverage thresholds (5-25 reads).

To rule out that the coding somatic variants that we observed AI on might in some cases alter the activity of a regulatory element overlapping the gene coding region, thereby causing imbalanced expression of itself, we removed mutations with the top 5% highest *Sei* scores. The correlations remained virtually unchanged (R = 0.51, 0.45, 0.05, for missense, ES and nonsense, respectively), supporting an independent epigenetic mechanism. The comparable ASCA–reg-AI correlations for ES and missense variants simply confirm that chromatin accessibility drives allelic transcription in principle independently of coding consequence. This provides a substrate for differential selection on the resulting AI: missense changes in bona-fide cancer genes could still be favoured or disfavoured, whereas synonymous changes would generally remain neutral. In practice, however, because of the fairly small number of tumor with ATAC-Seq available, most of our variants with matched ASCA data fall in passenger genes, leaving too few driver-gene observations to direct test for such selection effects on ASCA, which we acknowledge as a limitation.

To further evaluate the hypothesis that epigenetic influences during cancer evolution determine mRNA AI, as suggested above, we analyzed DNA methylation data. In particular, we considered all CpG sites near the TSS (see Supplemental Material and Methods for definition) jointly as a measure of gene activity levels, for each individual.

Given that positive reg-AI reflects a preferential expression of the mutated allele, we hypothesized that the promoter regions of these genes might show partial methylation, associated with differential expression between alleles. To test this, we compared variants with positive reg-AI to their methylation status, distinguishing between partially methylated (M-value > 30% and < 70% of gene’s percentiles), consistent with monoallelic expression, and unmethylated (M-value < 30% of gene’s percentiles) promoter, consistent with biallelic expression.

We observed significant enrichment of partial DNA methylation of the promoter-proximal region among variants with positive reg-AI in cognate TSGs (Figure S10B). This association was evident across all SNVs in cognate TSGs (OR = 1.80, 95% CI = 1.24 – 2.68, FDR = 6.6e-3) and was more pronounced for high aa-impact missense mutations (OR = 2.70, CI = 1.45 – 5.42, FDR = 9.3e-3). It was significantly stronger in TSGs compared to passenger genes (Breslow-Day test *p* = 5.1e-3), suggesting a tumor suppressor-associated effect with potential to generate cancer driver AI.

These findings suggest that in TSGs showing positive reg-AI, it is likely that the *wild-type* allele is being selectively silenced through promoter hypermethylation; as a limitation of this test, DNA methylation is from arrays, and thus cannot be phased with the somatic mutation on which the mRNA reg-AI was measured. This epigenetic modification could suppress the expression of the *wild-type* allele, thereby boosting the impact of the mutated allele, conferring a second hit (loss-of-function) effect in these TSGs, analogous to CNA deletions which remove the *wild-type* allele in TSGs(5).

Overall, these statistical links with epigenomic state of regulatory elements, explain a part of the reg-AI we observe in somatic variants in cancer. This supports the hypothesis that a mechanism analogous to imprinting, resulting from a epimutation, can underlie reg-AI, though we cannot rule out that unidentified *cis*-acting non-coding mutations also contribute.

## Discussion

In this study, we performed an in-depth analysis of the prevalence, mechanisms, and prognostic consequences of mRNA-level allelic imbalances of driver somatic mutations in cancer.

We estimated that the non-CNA causes of mRNA-AI, here united under the umbrella term “reg-AI” (based on the suggested roles on transcriptional and post-transcriptional regulation), explained about half (46.6%) of the total pan-cancer occurrences of AI of somatic mutations. This challenges the notion that somatic CNAs are responsible for the allelic expression imbalance in tumors(8, 15, 18, 19). Our results broadly align with recent studies of cancer-specific mRNA-level AI in 96 neuroblastomas(14), and a smaller-scale analysis of 9 breast, lung and head-and-neck cancers(69) [MIM: 275355], where 34%–65% of allele-specific mRNA expression was not explained by CNA or tumor heterogeneity. Along the same lines, in 43 glioblastoma stem cell samples, the genes showing tumor-specific mRNA AI did not cluster along the chromosomes, interpreted to mean segmental CNA were an unlikely cause(13). Next, in 91 TCGA breast cancer and matched normal samples, it was reported that 4.3% genes within a CNA have tumor-specific mRNA allele imbalance, however importantly that 1.8% of genes outside any CNAs (which is the majority of the genomic loci) also exhibited differential mRNA AI by the same criteria(9). Considered together, these recent studies and our contribution bring attention to mechanisms involving non-coding genetic variants and/or epigenomic changes in driving cancer-specific allele expression imbalances.

We stringently accounted for allele imbalances at the DNA level and sample purity, and the analysis comprehensively included exonic somatic SNVs across all driver genes and across >8000 tumors, extending previous smaller-scale, focussed cancer studies(8–13, 15, 19) to a pan-cancer setting. Of the high relevance cancer types, we observed a highest prevalence of reg-AI-affected variants in melanoma, lung, endometrial, stomach, kidney and colon, suggesting that these tumors commonly harbor epigenomic changes that drive cancer by interacting with driver mutations. At the other end of the spectrum, somewhat expectedly, ovarian cancer was among the more affected by AI generated through CNA, in accord with a very high prevalence of somatic CNA events in high-grade serous carcinomas (HGSC)(70, 71).

Similar to the CNA interactions with driver changes(2, 4, 5, 7), we demonstrated that the epigenetic AI is subject to somatic selection, with mRNA-level AI tilting in favor of the mutant allele in some driver genes; this confers a selective advantage and mimics LOH events resulting from CNA. We identified specific genes displaying significant differences in the non-CNA, reg-AI in 8 genes (22 at a more permissive FDR<25%), normalizing the impactful missense somatic mutations to a baseline of effectively-synonymous (ES) somatic mutations. The association of the missense mutations with positive mRNA-AI (i.e. imbalance in favor of the mutant allele is compatible with at least some of these alleles acting in a dominant-negative manner. Methodologically, our AI-dN/dES test considers both DNA and mRNA-level genetic variation, but unlike various prior tests for allele-specific expression, does not require RNA-Seq from tumor-matched healthy tissues from the same patient; because this, it can be applied to considerably larger datasets such as TCGA, boosting power.

Our test can identify cancer genes from AI of high impact somatic mutations in mRNA data; in addition to finding significant various TSGs and prominently *TP53*, and individual examples of OGs; the test further implicated additional genes such as *AMN*, *TTC32*, *ANKRD23*, *CYP4F3*, and *SBSPON* as putative driver (Table S2). These differences in overall mRNA-AI in selected cancer driver genes resulted from a largely mutually exclusive occurrence between CNA-AI and reg-AI, reinforcing the idea that non-CNA causes of mRNA allele expression imbalance are relevant to generating two-hit alterations that are selected for in cancer. Our approach is further supported by contemporaneous work describing the *RVdriver* method and tool(16), which leverages transcriptomic imbalance data to identify cancer driver genes. By analyzing RNA variant allele frequencies of missense mutations, they demonstrated that RNA approaches can complement traditional DNA-based methods for cancer gene discovery and moreover that they can distinguish driver from passenger mutations within established cancer genes (in analogy to our dN/dES test).

The mRNA-AI favoring the mutant allele in high aa-impact missense somatic driver gene mutations strongly predicts overall survival of cancer patients. We anticipate that there exist tens of additional driver genes where mRNA-AI is relevant to cancer aggressiveness, which will be clarified with additional statistical power from additional cohorts with larger sample sizes. The overall pan-cancer effect on survival was substantial for the overall mRNA-AI and specifically for reg-AI, compared to other known correlates of patient survival such as intratumor heterogeneity.

Next, we addressed the questions of the mechanisms underlying the observed non-CNA AI of driver mutations at mRNA level. While our analysis of AI association with somatic variants does not explicitly address heritable effects, such as that of imprinting, and of germline eQTL variants generating mRNA-AI, we infer these are likely not broadly relevant here, as imprinting does not affect many genes, and strong eQTLs do not affect many individuals(15, 72). Instead, we suggest that the epigenetic changes in the tumor that generate many reg-AI effects (including ones we identify to be under selection) are more alike to the “random monoallelic expression” effects(29–32): somatic epimutations which are stable across cell divisions. These would generate a source of epigenetic heterogeneity in tumors that we show can form epistatic interactions with driver mutations, by analogy to CNA, increasing tumor fitness.

As a caveat of our analysis, we cannot formally exclude the effects of distal somatic mutations e.g. in enhancers located in intergenic regions or in introns, in generating the non-CNA mRNA-AI with driver potential. However, the somatic mutations in or near promoters are presumably more likely to be impactful than in distal elements, yet we found that even promoter-proximal mutations can explain only a small fraction of the non-CNA mRNA-AI.

Similarly, it is possible albeit rare that the driver somatic mutation itself acts as a QTL and affects transcription of the target gene, e.g. by changing the local epigenomic environment and/or by promoting or blocking binding of a transcription factor. We used two modern machine learning predictors to suggest this is a rare occurrence: *Sei*(65) and *Puffin-D* tool(62). There may be individual examples where exonic mutations are predicted to have strong effects, particularly with 5’-most, TSS-proximal exonic mutations where *Puffin-D* predicts strong effects in some mutations (Figure S4).

In addition to direct effects of mutations on transcriptional activity, as an additional result we identify that non-CNA AI of somatic mutations results from splicing-altering coding region mutations with driver potential. Altered splicing registers as differential mRNA abundance downstream because it changes coverage with RNA-Seq reads and/or triggers NMD in cases of frameshifts or inclusion of poison cassette exons.

Our analysis employs RNA allele imbalance signal to supports the hypothesis that positive selection on synonymous mutations in OGs and TSGs operates through splicing mechanisms(60). Here we additionally observe that, similarly so, the selected missense mutations in cancer genes may result in splicing effects reflected in mRNA-AI.

Overall, splicing effects and promoter effects of mutations notwithstanding, we infer that mRNA-level AI changes are in most cases not downstream of somatic mutations nor CNA, but are rather more likely to results from independently occurring epigenetic changes. We observed examples thereof in allelic differences in chromatin accessibility (as ATAC-Seq), and in the DNA methylation near the promoter that associated with reg-AI; both readouts reflect the altered activity of enhancers or promoters. Future studies using long-read sequencing of tumors will enable the somatic mutation to be phased with the epigenetic change in question(34, 73), confirming whether the somatic driver mutation is indeed in *trans* with the inactivating promoter DNA methylation, and/or in *cis* with the activating, open chromatin state(74).

We would note that our models do not establish the temporal order between the mutation and the genetic or epigenetic change causing the AI; regardless of their ordering, their co-occurrence in a cancer driver gene could be under selection. The AI-causal changes may be local epigenome alteration but may occur due to trans-acting factors; indeed global epigenome remodelling is not uncommon in tumors(55–57) and could provide the second-hit event for several somatic driver mutations at once. This would generate driver events via mRNA expression imbalance by analogy to how arm-level/chromosome-level CNA are able to interact with various point mutations.

Overall, we find that impactful somatic mutations in driver genes often associate with allelic imbalances at the mRNA level. Positive selection favors the abundance of high impact mutations in mRNAs of at least 18 driver genes, including those where the allelic imbalance results from causes other than CNA. These and dozens of additional genes may predict detrimental prognostic outcomes in affected patients, presumably by reflecting epigenome instability that contributes to intratumoral heterogeneity and generates two-hit driver events via epimutations. We encourage for mRNA-level AI of cancerous mutations to be regarded not only as a prognostic marker, but also a prospective therapeutic target in the context of personalized medicine, prioritizing patients for pharmacological intervention, similarly as was proposed for DNA-level AI resulting from somatic copy-number changes(4).

## Data and code availability

The published article includes datasets generated or analyzed during this study, which are included as Tables S2-S3. Code generated for this analysis is provided at https://github.com/gpalou4/regAI

## Acknowledgements

We are grateful to Marcell Veiner for running the neural networks *Sei* and *spliceAI* on our somatic mutation data relevant to AI analysis. Further, we thank all members of the Genome Data Science lab for discussions. We used a large language model (Google Gemini 2.5 Pro) for language editing in parts of the manuscript text. G.P.-M. was funded by an AGAUR FI fellowship, and F.S. was funded by the ICREA Research Professor program. Work in the lab of F.S. was supported by an ERC StG “HYPER-INSIGHT” (757700), ERC CoG “STRUCTOMATIC” (101088342), Horizon2020 project “DECIDER” (965193), Spanish government project “REPAIRSCAPE”, CaixaResearch project “POTENT-IMMUNO” (HR22-00402), the SGR funding of the Catalan government, and the Severo Ochoa Centers of Excellence award of the Spanish government to the hosting institution. The results published here are in whole or part based on data generated by the TCGA Research Network (https://www.cancer.gov/tcga).

