## Supplementary Text S1-S3 for "Copy number-independent allelic imbalance in mRNA is selected in cancer and has prognostic relevance"

**Contents:**

**Text S1.** Post-transcriptional cause of reg-AI via splice-disrupting mutations in cancer driver genes – Tumor suppressor genes are often silenced by splice-disrupting missense variants – Splicing effects of exonic coding mutations are common in oncogenes and cause mRNA allelic imbalance.

**Text S2.** Somatic coding mutations rarely directly activate transcription in driver genes.

**Text S3.** Promoter-proximal non-coding mutations contribute to regulatory allelic imbalance in cancer driver genes.

### Text S1

*Post-transcriptional cause of reg-AI via splice-disrupting mutations in cancer driver genes*

The reg-AI signal in our data encompasses all types AI variation at the mRNA level (mRNA-AI) that cannot be explained by DNA-level AI originating from CNA (CNA-AI). Mechanisms that give rise to reg-AI in cancer genes are of interest. A prime example are somatic mutations altering splicing, which can have profound impacts on gene expression in cancer, for instance by inducing exon skipping, activating cryptic splice sites[1] and/or resulting in frameshifts that trigger nonsense-mediated mRNA decay (NMD)[2,3].

We therefore hypothesized that splicing changes might be a common post-transcriptional mechanism underlying reg-AI. To test this, we used two deep learning-based splicing predictors, *SpliceAI*[4] ($SP$) and *Pangolin*[5] ($PG$), to estimate the impact of somatic variants affecting splicing within a 50-bp window (see Methods). *SpliceAI* provides four delta score predictions—Donor Loss ($SP_{\mathrm{DL}}$), Donor Gain ($SP_{\mathrm{DG}}$), Acceptor Loss ($SP_{\mathrm{AL}}$), and Acceptor Gain ($SP_{\mathrm{AG}}$)— each quantifying the predicted change in splice site usage between the *mutated* and the *wild-type* alleles. *Pangolin* reports two analogous delta scores representing the maximum predicted increase ($PG_{\mathrm{gain}}$) or decrease ($PG_{\mathrm{loss}}$) in splice site activity. To capture the single strongest predicted effect per variant, we defined $SP_{\mathrm{Max\Delta}}$ as the largest absolute *SpliceAI* delta score across all four predictions and $PG_{\mathrm{Max\Delta}}$ as the largest absolute value of the two *Pangolin* scores.

We categorized somatic variants as splice-disrupting if their *SpliceAI* $SP_{\mathrm{Max\Delta}}$ score was ≥ 0.15 or their *Pangolin* $PG_{\mathrm{Max\Delta}}$ score was ≥ 0.1. Across all our dataset, 4.7% and 7.9% of total exonic variants were identified as splice-disrupting by $SP_{\mathrm{Max\Delta}}$ and $PG_{\mathrm{Max\Delta}}$, respectively. Stratifying variants by their association with reg-AI we found that 4.6-7.1% of variants associated with positive reg-AI were splice-disrupting, similar to 4.7-7.6% for variants showing no significant reg-AI. However, 10-15.4% of variants exhibiting negative reg-AI were predicted to be splice-disrupting. Among cognate TSGs, 4.9% of mutations with no significant reg-AI were splice-disrupting by *SpliceAI* (Fig. 2B), compared to 10.9% of those with significant reg-AI (OR = 2.4, $p$ = 4.1e-5, Fisher exact test; positive reg-AI: 11.5%; negative reg-AI: 10.1%). For cognate OGs we observed a similar pattern (4.1% no AI vs 10.2% reg-AI, OR = 2.7, $p$ = 1.3e-3; positive reg-AI: 11.8%; negative reg-AI: 7.8%). This indicates that exonic mutations that are predicted to alter splicing, considered as a set, tend to reduce the expression of the mutated allele; individual cases in the opposite direction are nonetheless possible, such as observed in cognate cancer genes.

As a prominent example, we examined three known splice-disrupting exonic driver variants in the *TP53* gene[6,7] (Figure S3A). Notably, the 3′ terminal nucleotide of *TP53* exon 4 (codon 125), harboring recurrent synonymous mutations, had a mean $SP_{\mathrm{DL}}$ delta score of 0.7 and $PG_{\mathrm{loss}}$ of 0.72, across the 3 possible somatic synonymous variants at this codon. These *TP53* mutations occur in a total of 32 tumor samples in our dataset, of which 7 exhibit a significant negative reg-AI. Similarly, codon 224 of *TP53* exon 6, showed a $SP_{\mathrm{DL}}$ of 0.9 and $PG_{\mathrm{loss}}$ of 0.78 in one synonymous variant (n = 3, of which 1 has negative reg-AI). Lastly, the codon 331 of *TP53* exon 9, exhibited a mean $SP_{\mathrm{DL}}$ of 0.62 and $PG_{\mathrm{loss}}$ of 0.85 among two low aa-impact missense variants (n = 5 occurrences, of which 1 has negative reg-AI), providing an example where a driver missense variant in *TP53* likely exerts its effect through splicing alteration, rather than through amino acid substitution.

**Tumor suppressor genes are often silenced by splice-disrupting missense variants**

The previously observed global enrichment of splice-disrupting variants among mutations exhibiting negative reg-AI (Fig. 2B) was anticipated because many splicing motif disturbances result in out-of-frame splicing products, which usually premature stops and trigger NMD[3,8].

To formally check this, we tested whether the proportion of predicted splice-disrupting variants was significantly higher for mutations associated with negative reg-AI (and separately for positive reg-AI) compared to those with no reg-AI (Figure S3B). This analysis was performed for distinct gene category (OGs, TSGs, or passenger genes as controls) and predicted functional impact of SNVs (all SNVs, high aa-impact missense or effectively synonymous; the latter also stratified further into low aa-impact missense and synonymous groups).

Within TSGs, a strong significant enrichment of splice-disrupting variants was observed for mutations leading to negative reg-AI among the ES variants ($SP_{\mathrm{Max}\Delta}$ OR = 3.61, FDR = 8.4e-3; $PG_{\mathrm{Max}\Delta}$ OR = 5.44, FDR = 1.3e-2, Figure S3B-C), but not within high aa-impact missense variants. This enrichment was significantly higher than that found in passenger genes ($SP_{\mathrm{Max}\Delta}$ and $PG_{\mathrm{Max}\Delta}$ meta-*p* = 2.2e-2, Breslow-Day test, Figure S3C), highlighting these splicing effects are selected during tumor evolution. The enrichment within TSGs was predominantly driven by variants predicted to cause splice donor loss ($SP_{\mathrm{DL}}$ OR = 8.26, FDR = 8.1e-4; $PG_{\mathrm{loss}}$ OR = 4.47, FDR = 3.8e-3).

Given that ES variants encompass both low aa-impact missense and synonymous variants, and considering that synonymous variants are known to modulate splicing patterns in cancer genes[3,6,8], we further checked whether the low aa-impact missense mutations within this ES group could similarly affect splicing and contribute to reg-AI. Indeed, these low aa-impact missense mutations showed a significant splicing effect on negative reg-AI ($SP_{\mathrm{Max}\Delta}$ OR = 9.34, FDR = 2.4e-2; $PG_{\mathrm{Max}\Delta}$ OR = 6.97, FDR = 6.2e-2), but not observed for synonymous mutations ($SP_{\mathrm{Max}\Delta}$ OR = 2.69, FDR = 0.22; $PG_{\mathrm{Max}\Delta}$ OR = 2.94, FDR = 0.16) (Figure S3B). In detail, 17.9% of low aa-impact missense mutations with negative reg-AI exhibited high splice-disrupting scores, in contrast to 6.9-9.3% ($SP_{\mathrm{Max}\Delta}$ and $PG_{\mathrm{Max}\Delta}$, respectively) in passenger genes, or 7.6-9.8% in TSGs in noncognate cancer types. This 8.6-11% difference compared to passenger genes quantifies the low aa-impact missense variants in TSGs that are under a selective pressure for splicing alterations ( $SP_{\mathrm{Max}\Delta}$ and $PG_{\mathrm{Max}\Delta}$ meta-*p* = 3.2e-2, Breslow-Day test, Figure S3C). For synonymous variants, the observed difference was 4.3-7% but not significant. Therefore, many missense somatic mutations drive cancer through splicing effects, rather than or in addition to alterations to the amino acid sequence, as was proposed for synonymous coding mutations[6].

Curiously, cognate TSGs as a group also had a significant enrichment of splice-disrupting variants (predicted by *SpliceAI*) among mutations with positive reg-AI ($SP_{\mathrm{Max}\Delta}$ OR = 3.27, FDR = 1.9e-3, Figure S3B-C) and it was significantly higher than that observed in passenger genes (*p* = 3e-5, Breslow-Day test, Figure S3C; of note, *Pangolin* was not included for meta-*p* calculation for this specific comparison because its predicted direction of effect did not align with that of *SpliceAI*). These positive reg-AI suggest dominant-negative acting TSG variants or isoforms.

Focusing on the *TP53* gene, it notably exhibited this enrichment of splice-disrupting variants among high aa-impact missense mutations with negative reg-AI ($SP_{\mathrm{Max}\Delta}$ OR = 6.81, FDR = 1.6e-2; $PG_{\mathrm{Max}\Delta}$ OR = 3.67, FDR = 5.2e-2) (Figure S3B). This was significantly greater than that found in passenger genes ($SP_{\mathrm{Max}\Delta}$ and $PG_{\mathrm{Max}\Delta}$ meta-*p* = 2.4e-2, Breslow-Day test, Figure S3C). These findings are consistent with prior work on exonic variants that disrupt splicing of *TP53*; previously, synonymous variants were reported[3,6,8], and here we suggest high aa-impact missense variants can have substantial splicing effects.

We further asked which other TSGs exhibit links between splicing altering variants and negative reg-AI at mRNA level (Figure S3D). Beyond *TP53*, no other individual TSG showed a significant enrichment for such variants at an FDR < 10%. However, we observed suggestive enrichments (FDR < 15%) for variants within low aa-impact mutations in two TSGs: *BAP1* (OR = 21.7, FDR = 0.13) and *CDH1* (OR = 19.7, FDR = 0.13).

### Splicing effects of exonic coding mutations are common in oncogenes and cause mRNA allelic imbalance

Next, we asked if the association to splicing effect scores was also coupled to mRNA allelic imbalance also in OGs. Interestingly, the effect on cognate OGs for ES variants was contrary to the trend in TSG: we identified an enrichment of splice-disrupting mutations among those exonic ES mutations having positive reg-AI ($SP_{\mathrm{Max}\Delta}$ OR = 6.55, FDR = 1e-2; $PG_{\mathrm{Max}\Delta}$ not significant, Figure S3B). This enrichment was significantly higher than that found in passenger genes ($SP_{\mathrm{Max}\Delta}$ and $PG_{\mathrm{Max}\Delta}$ meta-*p* = 1.8e-5, Breslow-Day test, Figure S3C). The enrichment within OGs was predominantly driven by variants predicted to cause splice acceptor loss ($SP_{\mathrm{AL}}$ OR = 14.3, FDR = 1.4e-3; $PG_{\mathrm{loss}}$ not significant).

This pattern indicates that splicing changes may enhance the expression of the mutated allele in cognate OGs (gain-of-function effect), in agreement with positive selection on synonymous mutations in OGs, proposed to act via splicing[6]. Further ES stratification showed that this enrichment is primarily stemmed from low aa-impact missense mutations ($SP_{\mathrm{Max}\Delta}$ OR = 8.13, FDR = 3.3e-3, Figure S3B-C). In contrast to the effect of ES mutations, no enrichment of splice-disrupting associated with positive reg-AI within high aa-impact missense mutations was found ($SP_{\mathrm{Max}\Delta}$ OR = 1.78, FDR = 0.7).

Next, in a gene-level analysis, we did not find oncogenes with a significant enrichment of positive reg-AI for ES or missense variants with splice-disrupting scores (Figure S3D).

In summary, coding regions variants can lead to an allele-specific reduction or increase in mRNA levels, by disrupting splicing. This can be exploited by cancer evolution: in TSGs we generally observed a decreased expression of the mutated allele via reg-AI, resulting in a loss-of-function. Conversely, our data suggests OGs can leverage these splicing alterations to exert a gain-of-function effect, characterized by an increased expression of the mutated allele.

### Text S2

<u>*Somatic coding mutations rarely directly activate transcription in driver genes*</u>

We investigated other mechanisms that give rise to reg-AI in cancer. We considered whether the somatic coding mutations associated with mRNA allelic imbalance might themselves modulate transcriptional activity. In principle, a somatic exonic mutation could alter the local epigenetic landscape—by perturbing nearby *cis*-regulatory elements (CREs) such as intragenic enhancers—or, if sufficiently proximal to the transcription start site (TSS), directly influence promoter function.

To test this hypothesis, we applied two deep-learning approaches: *Sei*[9] predicts chromatin state annotations (including active transcription chromatin states TN1-TN4, see Methods) from local DNA sequence, to assess how coding mutations shift chromatin toward a transcriptionally active state; *Puffin-D*[10] predicts the activity of gene promoters from DNA sequence within a 100kb-wide window, as GRO-Seq, PRO-Seq and CAGE scores. By comparing predicted scores for mutated versus reference sequences (the *ΔPuffin-D* score, see Methods), we assessed the direct impact of coding variants on promoter output.

We found that the *ΔPuffin-D* score of only 0.17% (for cancer genes) and 0.50% (for passenger genes) of the tested exonic variants exceeded the median of the positive control variants (*TERT* mutations known to affect promoter activity[11,12]) and only 0.60% and 1.53% was above the 1$^{st}$ (lower) quartile of positive controls (Fig. 2B). Thus, perhaps expectedly, exonic mutations are predicted to very rarely have large effects on transcriptional output. An interesting exception are exonic mutations at the very 5' ends of genes (first ~200 bp downstream of TSS), which can have large predicted effects (Figure S4). For example, individual mutations in six OGs (*CALR*, *GNAS*, *SF3B1*, *RAC1*, *BCL2L12*, *FUBP1*) and eight TSGs (*CDC73*, *ERCC4*, *EXT1*, *PPP6C*, *RPL5*, *SDHB*, *XPA*, *PPP2R1A*) yielded predicted *ΔPuffin-D* scores above the median of our positive controls. The median distance from these 24 exonic, expression-altering mutations, to the TSS was 18 bp. Importantly, only 16.67% (4/24) of these predicted promoter-activating mutations showed evidence of significant reg-AI in our data, compared to 5.92% for passenger genes (117/1976).

For exonic variants predicted to impact local chromatin activity by *Sei*, we classified variants with scores in the top 5% of our dataset as having potential transcriptional impact, given the absence of specific positive controls for comparison. When analyzing the relationship between these high-impact predictions and reg-AI across gene categories, we found that cognate TSGs showed a significant enrichment (Fig. 2B): 12% of SNVs with high-impact *Sei* scores exhibited positive reg-AI compared to only 5% of SNVs with no reg-AI (OR = 2.48, FDR = 1.95e-2). This enrichment was driven only by 23 SNVs from 7 TSGs, with the majority (n = 15) originating from *BCOR*, and the remainder distributed across *CDH1* (2), *FBXW7* (2), *CDKN2A* (1), *KEAP1* (1), *POLE* (1), and *SETD2* (1). Notably, 20 of these 23 variants were high aa-impact missense variants. In contrast, passenger genes showed the expected balanced distribution of approximately 5% of pos reg-AI versus 5% of no reg-AI (OR = 0.95), consistent with our top 5% threshold definition.

Overall, the direct transcriptional modulation associated with regulatory allelic imbalance can, albeit rarely, derive directly from somatic coding variants driving changes in promoter activity or local chromatin states. However, these rare examples do not explain the majority of reg-AI of somatic mutations in driver genes that we observed.

### Text S3

*Promoter-proximal non-coding mutations contribute to regulatory allelic imbalance in cancer driver genes*

While direct transcriptional effects of coding mutations proved rare, we considered whether reg-AI might instead result from non-coding somatic mutations (apart from CNA) affecting regulatory regions; this is analogous to germline eQTL (expression quantitative trait loci) variants often causing allele-specific expression by e.g. affecting transcription factor binding sites. A priori this explanation does not seem likely for our case of reg-AI selected cancer driver mutations, because driver somatic mutations affecting non-coding regulatory DNA are quite rare[13]. Known individual examples include *TERT* gene promoter driver mutations, and structural variants that translocate regulatory elements via promoter or enhancer hijacking[11,14].

We therefore considered the TCGA tumors from our original analyses that also had WGS data available (n = 777 tumors, from 22 cancer types; 44,373 WES variants available from those samples). We tested for co-occurring promoter-proximal mutations (conservatively, we include all variants ±1.5kb from the TSS, see Methods) in cancer driver genes with exonic variants exhibiting significant reg-AI (Figure S5).

Among these coding mutations, 55% of those with positive reg-AI (45 of 82) had co-occurring promoter-proximal non-coding mutations (either SNV or indel), compared to 44% for both no reg-AI (763 of 1,728) and negative reg-AI (21 of 55). In contrast, passenger genes showed the opposite pattern: 40% for negative reg-AI (482 of 1,202), 38% for no reg-AI (12,945 of 33,771), and 27% for positive reg-AI (186 of 697) (Figure S5). Thus, a promoter mutation co-occurs with a downstream exonic mutation exhibiting positive reg-AI 27% significantly more often in drivers than in passengers (55% vs 27%, OR = 3.33, $p$ = 6.1e-7) and 11% more often than in driver genes lacking reg-AI (55% vs 44%, OR = 1.54, $p$ = 6.8e-2). These excesses imply that ~11–28% of SNVs with positive reg-AI events in cancer driver genes can be attributed to nearby regulatory mutations, underscoring the role of TSS-proximal somatic variants as drivers of mRNA allelic imbalance in cancer.

Negative reg-AI values did not show significant differences between cancer and passenger genes, thus the selected effects of promoter-proximal mutations are only for overexpressing, but not for silencing an allele of a cancer driver gene.

We acknowledge some limitations of this analysis: first, that it is possible that somatic mutations in distal enhancers affect gene expression, however these are likely to have lower effects than the promoter-proximal mutations, which we tested above; second, because non-coding mutations are not phased with the corresponding coding mutations, we cannot determine whether their effects support or counter allelic imbalance. Nonetheless, the observed enrichment in cancer driver genes strongly suggests a meaningful association, even though the exact direction of the effect remains unclear.
