## Supplementary Figure S1-S10 for "Copy number-independent allelic imbalance in mRNA is selected in cancer and has prognostic relevance"

**Contents:**

**Figure S1.** Determinants of allelic imbalance (AI) in cancer genomes.
**Figure S2.** Proportion of SNVs with significant reg-AI versus CNA-AI among subclonal variants.
**Figure S3.** Association of splice-disrupting variants with decreased expression of mutated alleles.
**Figure S4.** Impact of coding mutations on transcription initiation predicted by the *Puffin-D* model.
**Figure S5.** Co-occurrence of promoter-proximal somatic mutations with exonic variants exhibiting regulatory allelic imbalance.
**Figure S6.** Allelic imbalance is subject to selection in cancer transcriptomes, potentially contributing to tumorigenesis.
**Figure S7.** CNA-driven versus regulatory allelic imbalance patterns in positively selected tumor suppressor genes.
**Figure S8.** CNA-driven versus regulatory allelic imbalance patterns in positively selected oncogenes.
**Figure S9.** Positive selection on regulatory allelic imbalance has a detrimental effect on overall cancer prognosis.
**Figure S10.** Suggestive evidence for an epigenetic cause in non-CNA allelic imbalance.

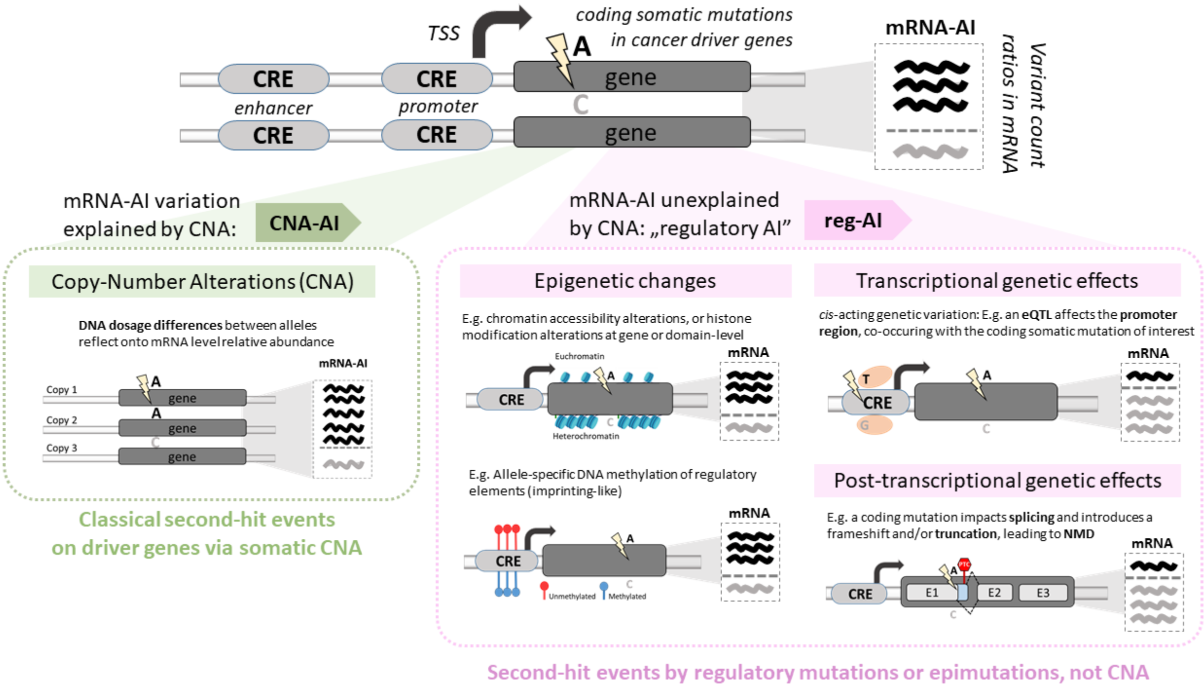


**Figure S1. Determinants of allelic imbalance (AI) in cancer genomes**

Illustration of two distinct paths through which AI may favor the expression of the mutant allele following an SNV event: (1) by copy number alterations (CNA) alone (left) or (2) by genetic or epigenetic variations (right). We attribute this latter, non-CNA AI, to epigenetic changes and/or to genetic variations that influence *cis*-regulatory elements (CREs), thereby affecting transcriptional regulation. We collectively refer to these non-CNA mechanisms as regulatory AI (reg-AI). Examples of epigenetic variation include histone modifications, chromatin accessibility or remodeling changes that occur without an underlying genetic variant (epimutation), as well as allele-specific DNA methylation, analogous to imprinting events. In some cases, a coding mutation can overlap with an intragenic CRE (e.g. an enhancer), disrupting a transcription factor (TF) binding site and producing downstream epigenetic effects. Alternatively, coding mutations may interfere with splicing or result in a nonsense mutation, both of which can introduce a premature termination codon (PTC), triggering nonsense-mediated decay (NMD).

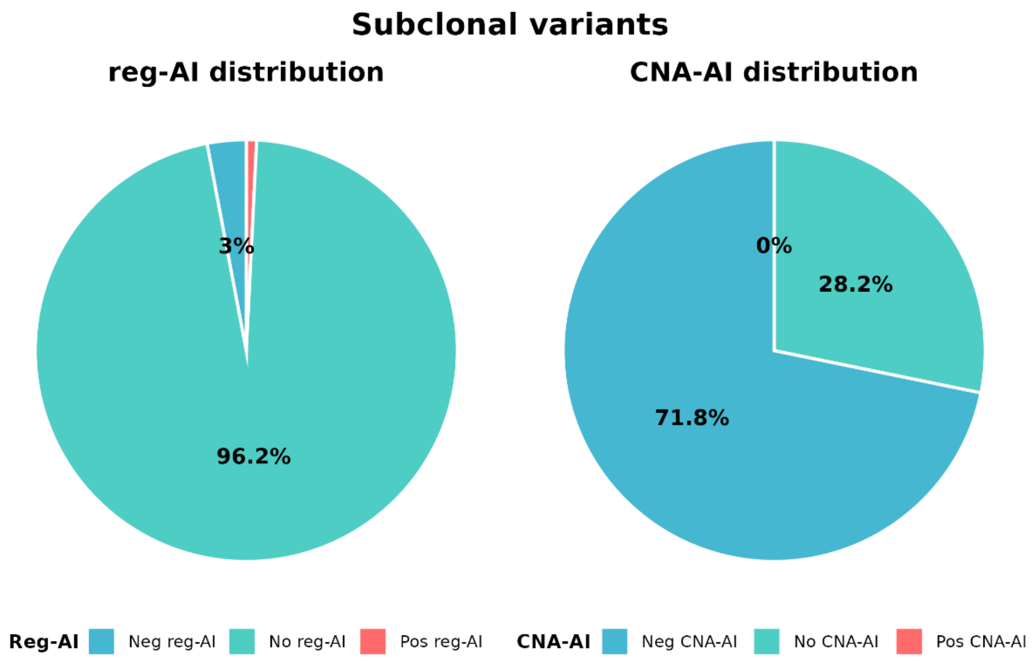


**Figure S2. Proportion of SNVs with significant reg-AI versus CNA-AI among subclonal variants.**

Pie chart displaying the distribution of allelic imbalance significance categories among putatively subclonal variants from cancer driver genes. The analysis includes 638 somatic SNVs (from a total of 41,631 variants – 1.5% – in cancer genes) meeting the following criteria: subclonal status $\left(\text{defined by } \frac{DNA\_VAF_{obs}}{purity} < 0.2\right)$, significant mRNA-AI, located in cancer genes (excluding passenger genes), excluding variants with deletions of the mutated allele ($ACN_{\mathrm{MUT}} = 0$), and excluding nonsense mutations. Each variant is categorized by its significance status for reg-AI and CNA-AI separately, with pie segments representing the proportion of variants showing significant positive or negative reg-AI, significant positive or negative CNA-AI, or no significance for either test.

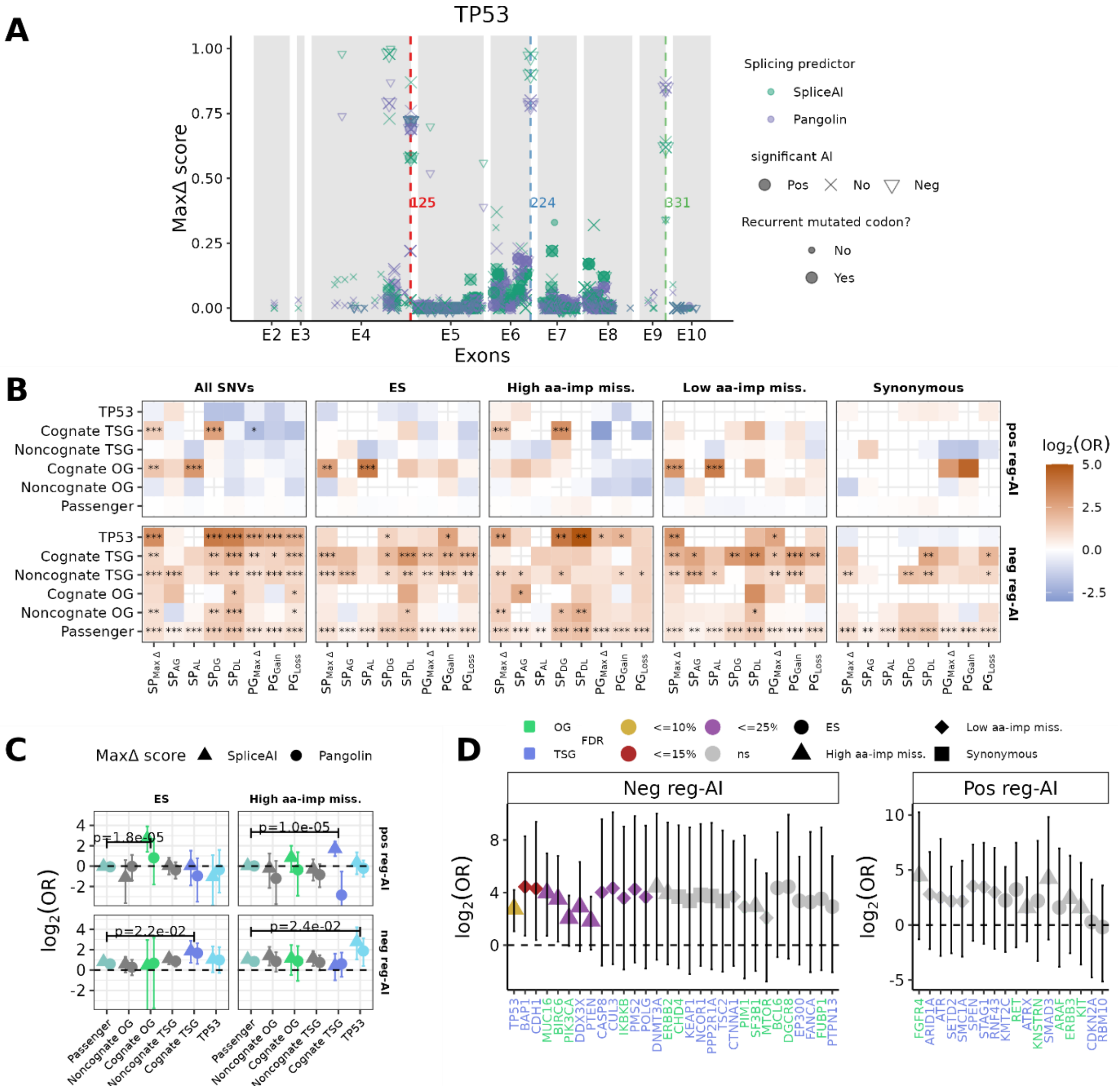


**Figure S3. Association of splice-disrupting variants with decreased expression of mutated alleles.**

**A,** Distribution of *SpliceAI*[1] ($SP_{\text{Max}\Delta}$) and *Pangolin*[2] ($PG_{\text{Max}\Delta}$) MaxΔ-scores for all exonic somatic single nucleotide variants (SNVs) within the *TP53* gene, plotted according to their sorted genomic coordinates (displaying exonic regions only). We defined $SP_{\text{Max}\Delta}$ as the largest absolute *SpliceAI* score across all four predictions and $PG_{\text{Max}\Delta}$ as the largest absolute value of the two *Pangolin* scores (see Methods). Individual SNVs are categorized based on their regulatory allelic imbalance (reg-AI) status: positive reg-AI, no reg-AI, or negative reg-AI. Note: While a specific SNV has a single *SpliceAI/Pangolin* score, its reg-AI status can vary across different patient samples. SNVs that are recurrently mutated at the same codon (observed in ≥ 5 samples within this dataset) are highlighted by larger point sizes. Horizontal dashed lines indicate the positions of three well-characterized splice-disrupting synonymous driver variants in *TP53*: codon 125, codon 224, and codon 331[3]. **B**, Enrichment analysis of predicted splice-disrupting mutations ($SP_{\text{Max}\Delta}$ ≥ 0.15 for *SpliceAI*; $PG_{\text{Max}\Delta}$ ≥ 0.10 for *Pangolin*) among those exhibiting either positive or negative reg-AI (preferential expression of the *mutated* or *wild-type* allele, respectively). The analysis is stratified by variant functional impact based on *VARITY_ER*[4,5] (see Methods): all SNVs, high or low aa-impact missense, synonymous-only and effectively synonymous (ES) which includes low aa-impact missense and synonymous grouped. It is also stratified by gene classification: noncognate and cognate oncogenes –OGs–, noncognate and cognate tumor suppressor genes –TSGs–, *TP53,* and passenger genes as controls. "Cognate" means SNVs matched to the cancer type where they are known to be a driver based on MutPanning[6]. Enrichments are evaluated for all five *SpliceAI* delta scores: Acceptor Loss ($SP_{\mathrm{AL}}$), Acceptor Gain ($SP_{\mathrm{AG}}$), Donor Loss ($SP_{\mathrm{DL}}$), Donor Gain ($SP_{\mathrm{DG}}$) and $\mathrm{Max}\Delta$ ($SP_{\mathrm{Max}\Delta}$), the latter representing the maximum value among the four categories ($SP_{\mathrm{AG}}$, $SP_{\mathrm{AL}}$, $SP_{\mathrm{DG}}$, $SP_{\mathrm{DL}}$) for each mutation. Similarly, we show the enrichments for the three *Pangolin* delta scores: Gain ($PG_{\mathrm{Gain}}$), Loss ($PG_{\mathrm{Loss}}$) and $\mathrm{Max}\Delta$ ($PG_{\mathrm{Max}\Delta}$). Odds Ratios (ORs) were calculated using the Fisher Exact test and presented in a $\log_2$ scale, color-coded from low (blue) to high (red) values. False discovery rates (FDRs) are indicated as follows: *** FDR < 0.001, ** FDR < 0.05 and * for FDR < 0.15. **C**, Enrichment analysis as in B, but displaying ORs and 90% Confidence Intervals (CI) for splice-disrupting mutations defined by *SpliceAI* ($SP_{\mathrm{Max}\Delta}$ ≥ 0.15) and *Pangolin* ($PG_{\mathrm{Max}\Delta}$ ≥ 0.10) scores. Enrichments within each gene category are compared against the passenger genes ORs as a baseline using Breslow-Day test, assessing homogeneity of ORs. Only significant *p*-values (< 0.05, meta-*p* from $SP_{\mathrm{Max}\Delta}$ and $PG_{\mathrm{Max}\Delta}$) from the test are shown. **D**, Similar to panel C, an enrichment analysis was performed on individual cancer genes (only TSGs and OGs), separately for positive and negative reg-AI. The results are displayed as $\log_2$ OR (Y-axis) with its 90% CI, highlighting the ranking of top 30 genes that were significant (FDR < 10%) or potential candidates (FDR < 15% or 25%). Enrichments were performed stratifying by SNV functional impact (ES, high or low aa-impact missense variant, and synonymous-only) and by splice-disrupting prediction scores from both *Pangolin* ($PG_{\mathrm{Max}\Delta}$) and *SpliceAI* ($SP_{\mathrm{Max}\Delta}$), provided there was a sufficient number of mutations. For each gene, and among the eight total combinations, the single most significant result (lowest FDR) was chosen to display.

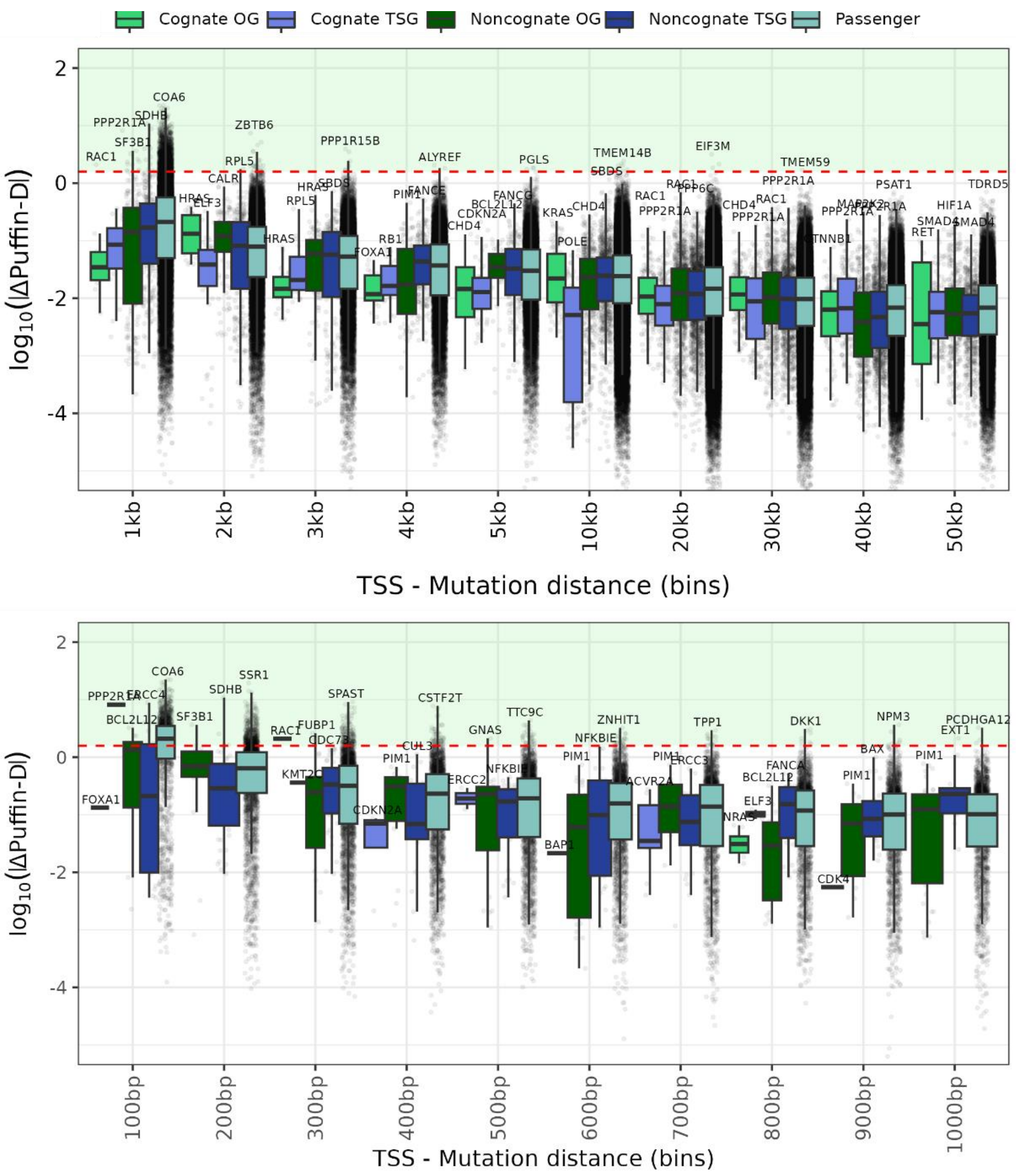


**Figure S4. Impact of coding mutations on transcription initiation predicted by the *Puffin-D* model.**

The *Puffin-D*[7] neural network was used to predict changes in gene promoter activity (measured by GRO-Seq, PRO-Seq, and CAGE scores) based on a DNA sequence within a 100kb window around the somatic mutation (see Methods). We analyzed the $\log_{10}$ of the absolute change in Δ*Puffin-D* scores (Y-axis) when exonic mutations were introduced in this sequence window, comparing mutated sequences to the reference DNA sequence. For this, we integrated the predictions over a 100nt segment centered on the gene's transcription start site (TSS). Mutations were grouped by genomic distance from the TSS in different intervals (top panel: ranging from 1kb to 50kb; bottom panel: ranging from 100bp to 1000bp) and classified by gene type, including cognate and noncognate TSGs and OGs, and passenger genes. To provide a benchmark for relevant effect sizes (green box), we included a threshold based on known somatic mutations affecting the promoter of *TERT* gene[8,9] as a positive control (horizontal dashed line). We also highlight the top gene with highest Δ*Puffin-D* score for each gene category and bin. While exonic mutations are generally predicted to have minimal effects on transcriptional output, a notable exception is observed at the very 5' ends of genes (within the first 200bp post-TSS), where exonic mutations can lead to substantial predicted effects on transcription.

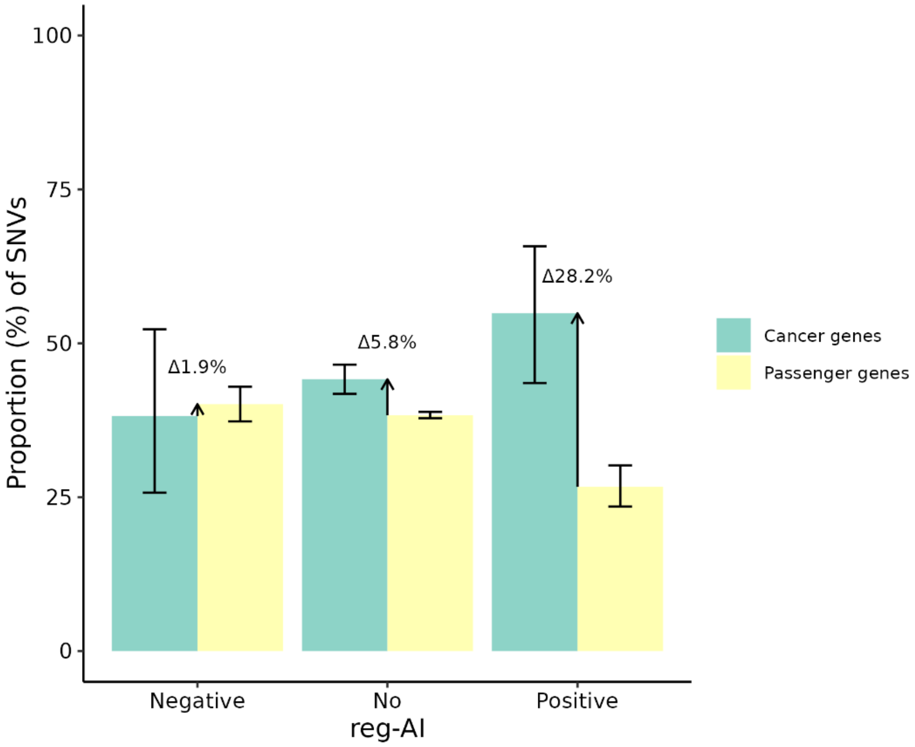


**Figure S5. Co-occurrence of promoter-proximal somatic mutations with exonic variants exhibiting regulatory allelic imbalance.**

Proportion of tumors harboring both promoter-proximal somatic mutations (SNVs or indels within ±1.5 kb of the transcription start site, see Methods) and exonic variants with regulatory allelic imbalance (reg-AI) in the same gene. Analysis includes TCGA tumors with whole-genome sequencing data available (n=777 tumors, 22 cancer types). Exonic variants are stratified by reg-AI status: positive reg-AI (preferential expression of the *mutant* allele), negative reg-AI (preferential expression of the *wild-type* allele), or no significant reg-AI. Genes are categorized as cancer drivers (all TSGs and OGs) or passenger genes. Red arrows indicate the magnitude of enrichment in cancer genes relative to passenger genes, suggesting positive selection for co-occurring regulatory mutations that enhance mutant allele expression in cancer drivers.

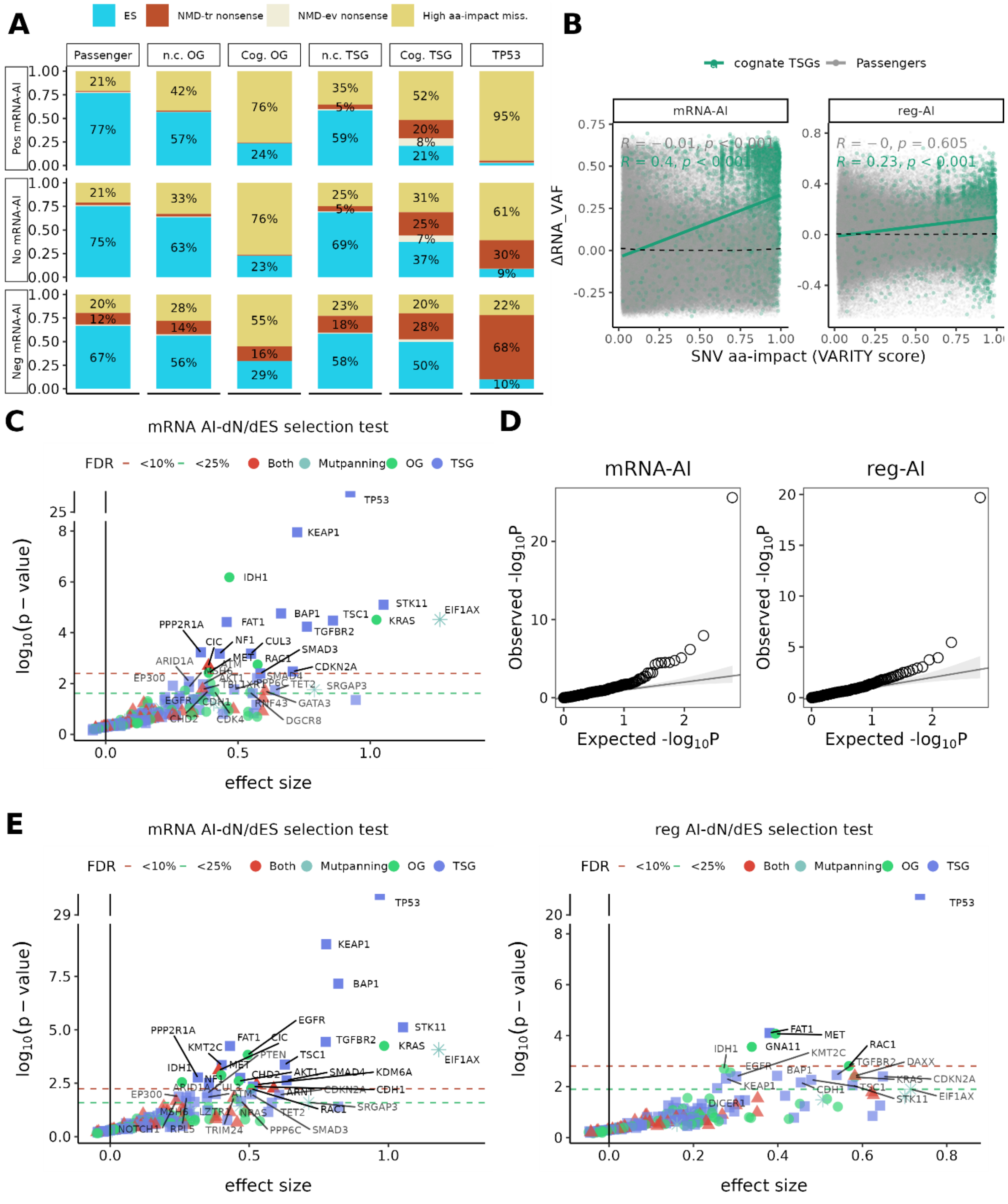


**Figure S6. Allelic imbalance is subject to selection in cancer transcriptomes, potentially contributing to tumorigenesis**

**A,** Relative frequencies of somatic SNVs, categorized by their predicted functional impact: effectively synonymous (ES), high aa-impact missense and nonsense. Nonsense mutations were further subdivided into those predicted to trigger nonsense-mediated mRNA decay (NMD-tr) and those predicted to evade (NMD-ev) based on known genomic rules[10] (see Methods). Proportions were further stratified by gene categories: passenger genes, oncogenes (OGs), tumor suppressor genes (TSGs) and *TP53* (analyzed independently). OGs and TSGs were additionally split into cognate (Cog.) and noncognate (n.c.). "Cognate" means SNVs matched to the cancer type where they are known to be a driver based on MutPanning[6]. Within each gene and functional impact category, proportions are grouped based on their mRNA-AI significance: no mRNA-AI (indicating no significant allelic imbalance at the mRNA level attributable to both CNA and regulatory effects), positive mRNA-AI (indicating preferential expression of the *mutated* allele) and negative mRNA-AI (indicating preferential expression of the *wild-type* allele). A higher proportion of positive mRNA-AI among missense mutations is indicative of selection pressure favouring increased mRNA expression of these *mutant* alleles. **B,** Relationship between SNV functional impact (by *VARITY*[4,5] scores, x-axis) and allelic imbalance strength ($\Delta RNA\_VAF$, y-axis) for missense mutations in cognate TSGs and passenger genes. Left panel shows mRNA-AI; right panel shows reg-AI. $\Delta RNA\_VAF$ represents residuals from beta-binomial models, calculated as the difference between observed and predicted RNA variant allele frequencies ($RNA\_VAF_{\mathrm{obs}} - RNA\_VAF_{\mathrm{pred}}$). Higher *VARITY* scores indicate greater predicted functional impact on amino-acid structure and function. Pearson correlation coefficients (R) and *p*-values are displayed for each gene category. The positive correlation in TSGs suggests that more functionally damaging missense mutations are associated with stronger mRNA-level AI. **C**, Volcano plot illustrating the results of our gene-level mRNA AI-dN/dES test for 356 total genes: OGs, TSGs, genes classified as both TSG/OGs, and MutPanning-only genes. The test adapts our pan-cancer beta-binomial framework, originally for detecting significant AI at the single SNV-level, to assess gene-level mRNA-AI (see Methods). The X-axis displays the effect size of the regression coefficient for the $SNV_{type}$ covariate (high aa-impact missense versus ES mutations) from the model. The Y-axis represents the raw upper-tail *p*-values from this regression, presented on a $-log_{10}$ scale. Genes exhibiting a statistically significant effect (FDR < 10%) or candidates (FDR < 25%) are highlighted, with its corresponding thresholds as horizontal dashed lines. A positive and statistically significant effect size for a gene indicates that its high aa-impact missense mutations are associated with a greater enrichment of the *mutant* allele's mRNA expression compared to the neutral baseline established by its ES mutations. This signature suggests that the observed AI itself is under positive selection in these identified driver genes, potentially contributing to their oncogenic role. **D**, Q-Q plots that compare observed versus expected $-log_{10}$ *p*-values from the AI-dN/dES gene-level tests for mRNA-AI (left), reg-AI (right). This helps to evaluate the extent of deviation from expected results under the null hypothesis. The lambda inflation factor (λ) shows minimal deviation for reg-AI (λ = 1.37) and moderate for mRNA-AI (λ = 1.63). **E**, Sensitivity analysis excluding subclonal variants. Replication of the AI-dN/dES analysis for mRNA-AI (left) and reg-AI (right) after excluding subclonal variants (using a heuristic approach, defined as $\frac{DNA\_VAF_{obs}}{purity} < 0.2$). Subclonal variants are mutations present in only a subset of tumor cells and can potentially bias selection estimates due to incomplete cellular representation. The modest reduction in detected genes −results remained qualitatively consistent with the original analysis at FDR<25%− reflects decreased power rather than systematic confounding by subclonal variants.

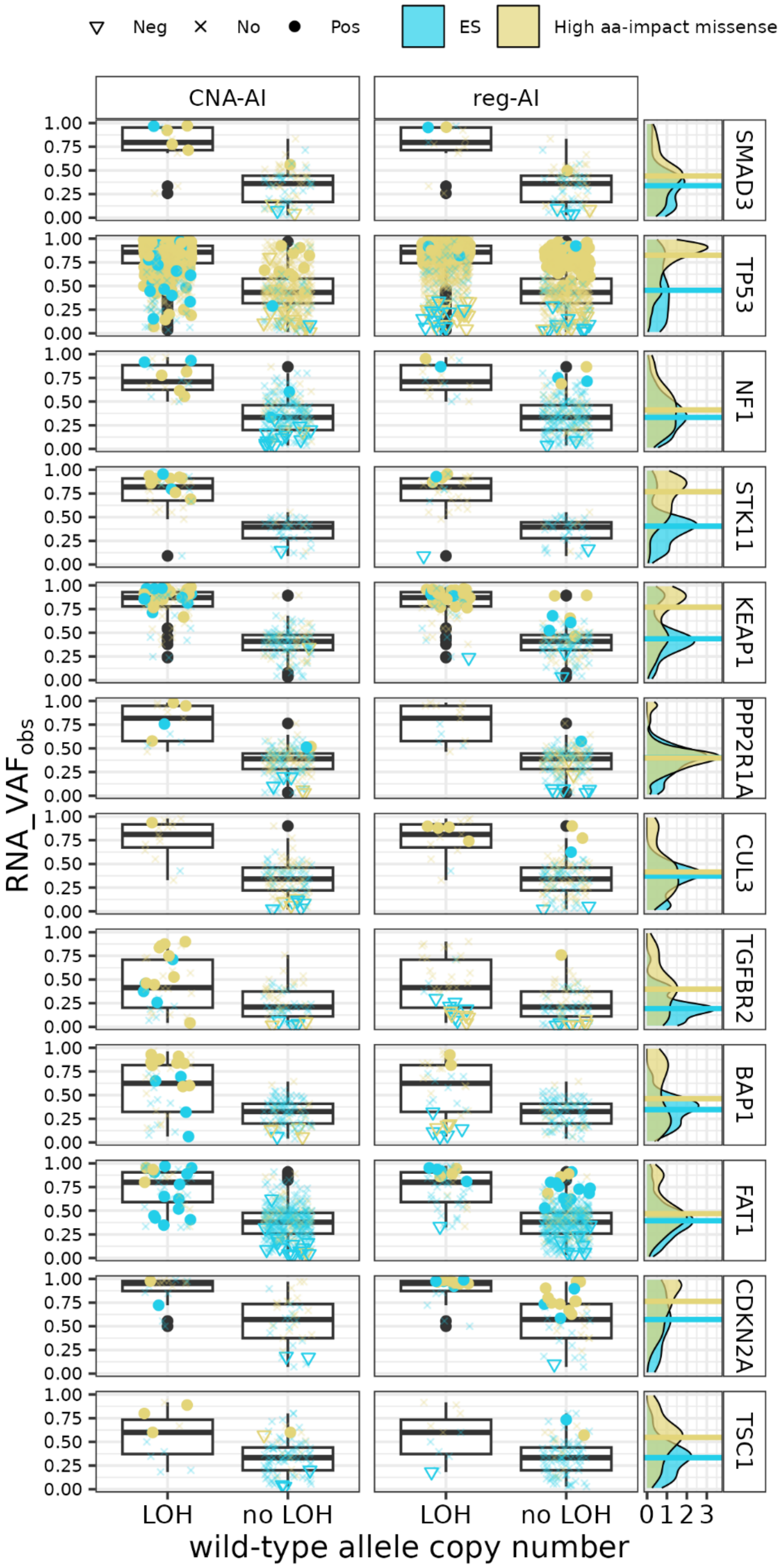


**Figure S7. CNA-driven versus regulatory allelic imbalance patterns in positively selected tumor suppressor genes.**

Distribution of $RNA\_VAF_{obs}$ for mutations plotted against ASCAT-assigned[11] *wild-type* allele copy number ($ACN_{WT}$, see Methods) for 12 positively selected TSGs identified by our mRNA AI-dN/dES test

(FDR < 25%). Variants are stratified by allelic imbalance significance: positive, negative, or non-significant for both CNA-AI and reg-AI. $ACN_{\mathrm{WT}}$ is used to create two categories: LOH ($ACN_{\mathrm{WT}}$ = 0) or no LOH (includes copy-neutral, $ACN_{\mathrm{WT}}$ = 1 and $ACN_{\mathrm{MUT}}$ = 1, and amplified, $ACN_{\mathrm{WT}} > 1$, SNV events). Variants are colored by functional impact: high amino acid (aa)-impact missense mutations (yellow) versus effectively synonymous variants (blue). The rightmost density plots show also median $RNA\_VAF_{\mathrm{obs}}$ values, confirming that high aa-impact missense mutations consistently exhibit higher expression than ES variants, supporting their identification as positively selected in our mRNA AI-dN/dES analysis. Expectedly, positive CNA-AI significant variants are enriched in LOH events, reflecting copy number-driven allelic imbalance of the *mutant* allele. In contrast, reg-AI significant variants are distributed equally across the two groups, demonstrating copy number-independent regulatory mechanisms. Individual variants may show significance for either CNA-AI or reg-AI, but rarely both simultaneously. For instance, *TP53* exhibits both reg-AI and CNA-AI significant variants, indicating dual mechanistic contributions. *BAP1*, *STK11*, and *KEAP1* show predominantly CNA-AI significance with limited reg-AI events. *CDKN2A*, *FAT1*, and *CUL3* display primarily reg-AI significant variants, suggesting regulatory-driven allelic imbalance.

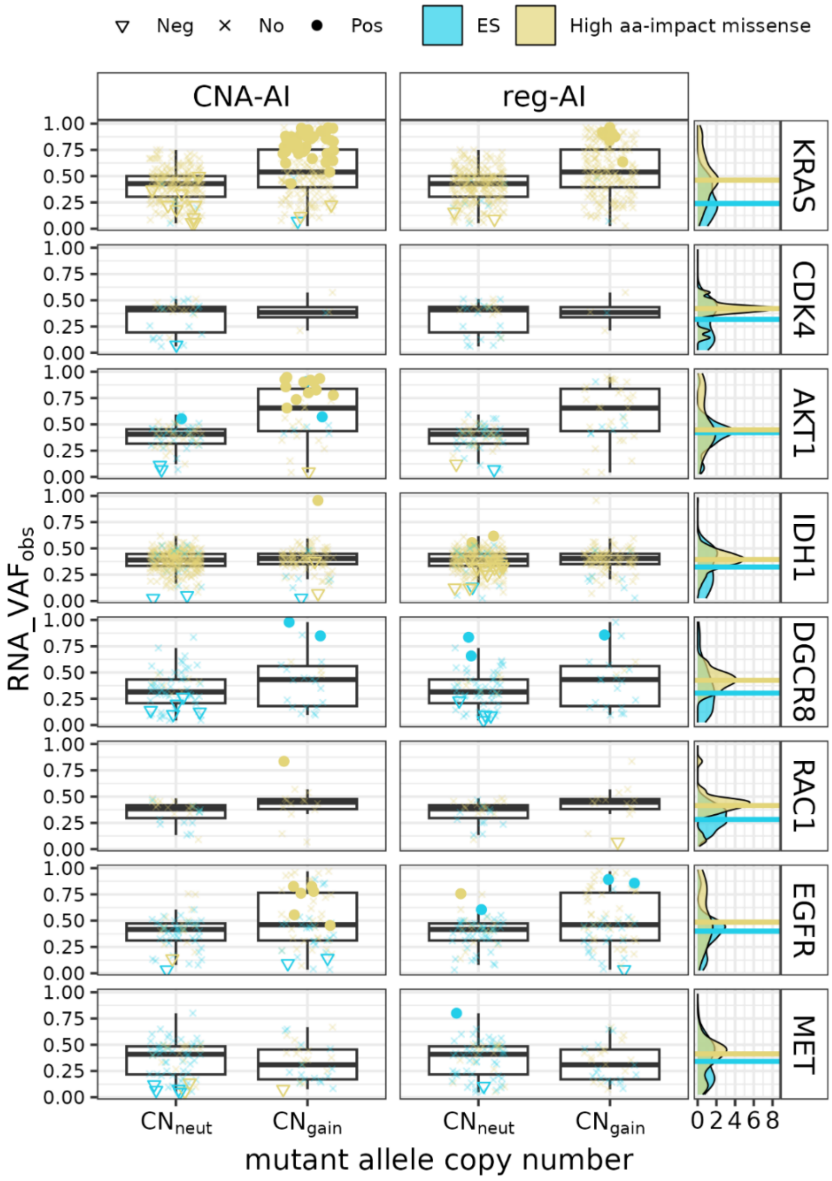


**Figure S8. CNA-driven versus regulatory allelic imbalance patterns in positively selected oncogenes.**

Distribution of $RNA\_VAF_{\mathrm{obs}}$ for SNVs plotted against ASCAT-assigned[11] *mutant* allele copy number ($ACN_{\mathrm{MUT}}$, see Methods) for 8 positively selected oncogenes identified by our mRNA AI-dN/dES test (FDR < 25%). Variants are stratified by allelic imbalance significance: positive, negative, or non-significant for both CNA-AI and reg-AI. $ACN_{\mathrm{MUT}}$ is used to create two categories: copy-number neutral ($CN_{\mathrm{neut}}$: $ACN_{\mathrm{WT}}$ = 1 and $ACN_{\mathrm{MUT}}$ = 1), or copy-number gained ($CN_{\mathrm{gain}}$: $ACN_{\mathrm{MUT}} > 1$); copy-number deletion cases were excluded ($ACN_{\mathrm{MUT}}$ = 0). Variants are colored by functional impact: high amino acid (aa)-impact missense mutations (yellow) versus effectively synonymous variants (blue). The rightmost density plots show also median $RNA - VAF_{\mathrm{obs}}$ values, confirming that high aa-impact missense mutations consistently exhibit higher expression than ES variants, supporting their identification as positively selected in our mRNA AI-dN/dES analysis. Expectedly, positive CNA-AI significant variants are enriched in amplification events, reflecting copy number-driven allelic imbalance of the *mutant* allele. Individual variants may show significance for either CNA-AI or reg-AI, but rarely both simultaneously. *KRAS and EGFR* exhibits both reg-AI and CNA-AI significant variants, indicating dual mechanistic contributions. *AKT1*, show predominantly CNA-AI significance with limited reg-AI events. *IDH1* display some reg-AI significant variants, suggesting regulatory-driven allelic imbalance. However, in general, there are very few reg-AI significant variants in our set of selected oncogenes.

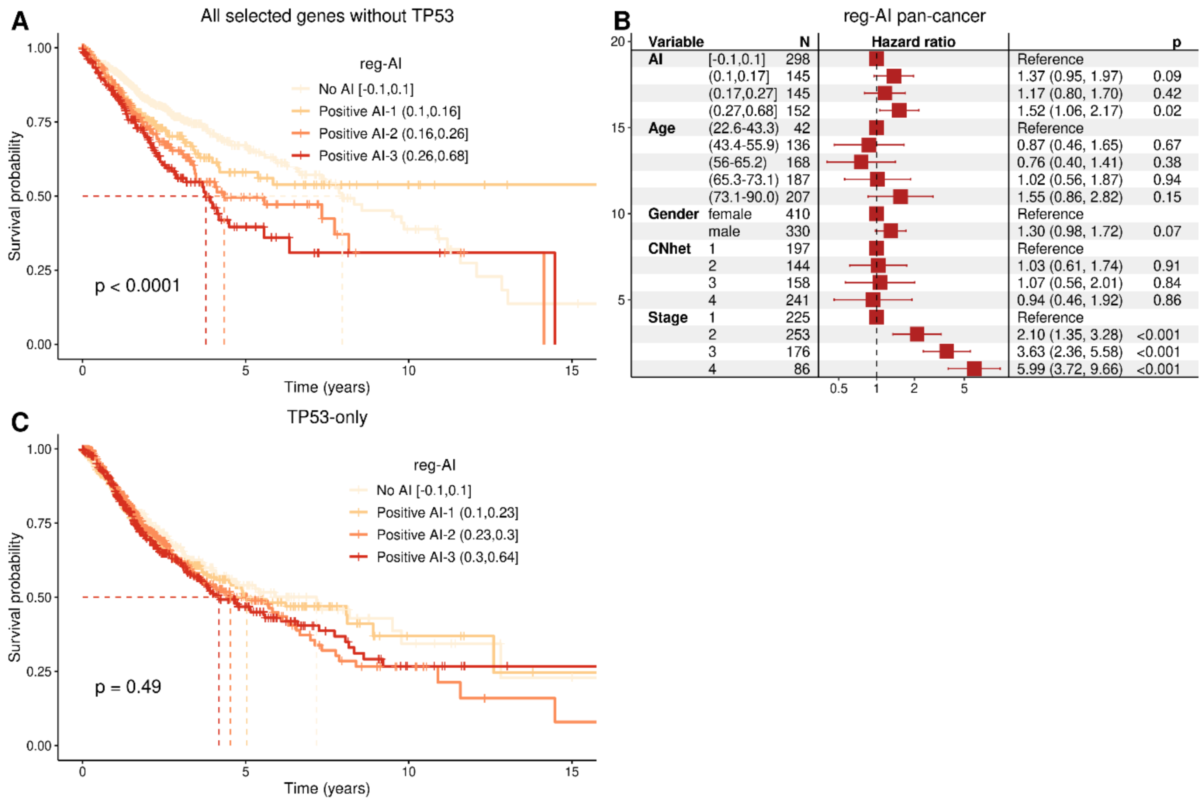


**Figure S9. Positive selection on regulatory allelic imbalance has a detrimental effect on overall cancer prognosis.**

**A**, Kaplan-Meier (KM) overall survival curves for pan-cancer patients harbouring at least one high aa-impact missense mutation in 22 genes under positive reg-AI selection (FDR < 25%; *TP53* excluded; ES mutations excluded). Patients are stratified by per-sample reg-AI score into three "positive" quantile bins groups ((0.1, 0.16], (0.16, 0.26], or (0.26, 0.68]) versus "neutral/reference" ([–0.1, 0.1]). This reg-AI score is calculated based on the $\Delta RNA\_VAF$, calculated as the difference between the $RNA\_VAF_{obs}$ and the $RNA\_VAF_{pred}$ by our beta-binomial model for each qualifying SNV (see Methods). The mean of these $\Delta RNA\_VAF$ across all considered SNVs within a given tumor was then used as a continuous reg-AI score for that sample. The indicated *p*-value is derived from a log-rank test. **B,** Cox proportional hazards survival analysis, showing the estimated hazard ratios (HR) for each variable, along with the 95% CIs and *p*-values. The results demonstrate a significant positive association between increasing reg-AI intensity and higher HRs. The model incorporated the following covariates: patient age (discretized into quintiles), sex, tumor stage, cancer type, intratumor genetic heterogeneity (estimated as copy-number heterogeneity, CNH)[12], tumor sample purity, and total sample CNA burden. Some of these covariates are excluded from the plot for better visualization. **C**, KM curves for pan-cancer overall survival, analogous to the analysis presented in panel A for reg-AI, but exclusively considering patients with high aa-impact missense mutations in *TP53*.

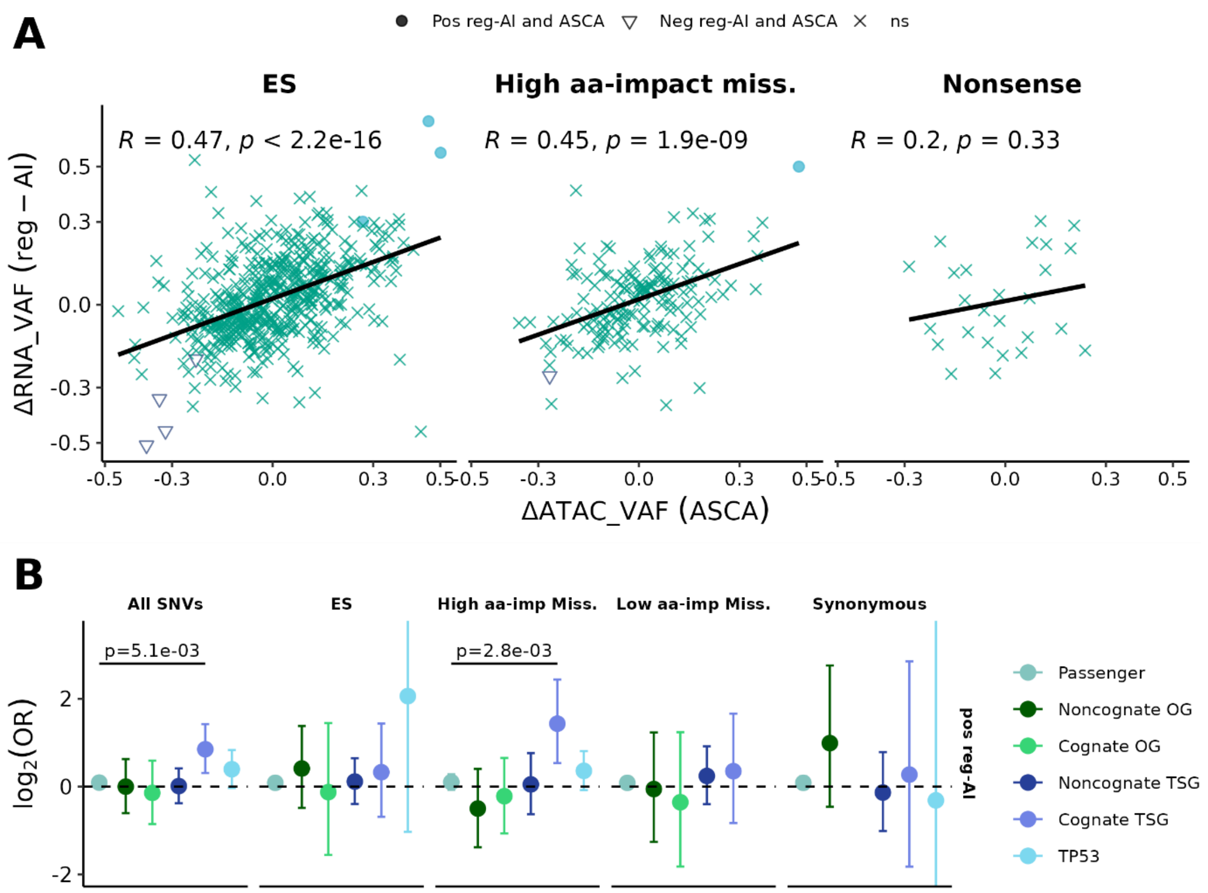


**Figure S10. Suggestive evidence for an epigenetic cause in non-CNA allelic imbalance.**

**A**, Chromatin accessibility correlates with regulatory allelic imbalance. Linear regression examining the relationship between regulatory allelic imbalance (reg-AI) and allele-specific chromatin accessibility (ASCA) across somatic mutation types. The y-axis shows $\Delta RNA\_VAF$ (residuals from the beta-binomial reg-AI model calculated as difference between $RNA\_VAF_{\mathrm{obs}}$ and $RNA\_VAF_{\mathrm{pred}}$), while the x-axis shows $\Delta ATAC\_VAF$ values (residuals from the beta-binomial ASCA model calculated as difference between $ATAC\_VAF_{\mathrm{obs}}$ and $ATAC\_VAF_{\mathrm{pred}}$). Analysis includes 782 variants from 381 TCGA patients with matched RNA-Seq and ATAC-Seq data (coverage threshold: ≥20 counts for both RNA-Seq and ATAC-Seq). Variants are stratified by functional impact: effectively synonymous (ES, left), high amino acid (aa)-impact missense (middle), and nonsense mutations (right). Points are shaped by allelic imbalance significance status, with Pearson correlation coefficients (R) and *p*-values displayed for each mutation category. **B**, DNA methylation status associates with regulatory allelic imbalance. Enrichment analysis comparing the frequency of positive reg-AI in genes with partial methylation versus unmethylated status within promoter-proximal regions (TSS200-1500 and first 2kb of gene bodies, see Methods). Results are stratified by gene categories (cognate and noncognate TSGs, OGs and passenger genes) and SNV functional impact (all SNVs, ES, high or low aa-impact missense, synonymous-only). $\log_2$ ORs with 95% CIs are estimated using Fisher's exact tests. Enrichments within each gene category are compared against the passenger genes ORs as a baseline using Breslow-Day test, assessing homogeneity of ORs. Only significant *p*-values (< 0.05) from the test are shown.
