## Supplementary Methods for "Copy number-independent allelic imbalance in mRNA is selected in cancer and has prognostic relevance"

**Contents:**

Comparison of AI contribution estimates with prior studies

Allele-specific chromatin accessibility (ASCA) analysis

Gene-level DNA methylation status

Variant effect predictions on chromatin profiles by Sei

Variant effect predictions on transcription initiation by Puffin-D

Variant effect predictions on splicing and enrichment analysis

Classification of nonsense mutations into NMD-triggering and NMD-evading

### Supplemental Material and Methods

*Comparison of AI contribution estimates with prior studies*

To replicate the findings from the Correia *et. al.* study[1] on reg-AI contribution in BRCA patients and compare our findings to theirs, we adjusted our dataset using a similar filtering approach. We applied more stringent filtering and then estimated our reg-AI and CNA-AI contributions as described in results section. These filterings included: (1) omitted adjustment by sample tumor purity in our beta-binomial models; (2) restricted the analysis to missense mutations; (3) focused exclusively on BRCA patients; (4) applying an RNA-Seq coverage threshold of at least 30 counts (sum of WT and MUT alleles); (5) focusing solely in the *PIK3CA* gene, all as per Correia *et. al.*[1]. Under these conditions, we found that our reg-AI contribution decreased from our original 46.6% to 16.7%, which closely aligns with the 16% reported by Correia *et. al.*[1]. Therefore we infer the difference stems from a more comprehensive analysis in our case, including various cancer types and genes and variants, and purity adjustment.

The PCAWG consortium performed a transcriptomic analysis of 1,188 RNA-Seq samples from several cancer types[2] using a logistic regression model to identify factors contributing to allelic imbalance. Their analysis attributed 86.1% of the explained total effect size to CNA. However, this approach relied on explicitly defined putatively causal variables in a model, estimated the contributions of the remaining tested variables (apart from CNA) on AI to be of 15.7%. Crucially, this type of modeling can estimate the relative contribution of non-CNA-AI (reg-AI) only due to the known variables that were quantified and included in the model, but ignores the non-CNA-AI (reg-AI) stemming from unknown sources. This underestimates the relative contribution of the non-CNA-AI, because it is likely an aggregate of multiple mechanisms, which may or may not have been measured and included in the model, for instance the cancer epigenetic evolution. In contrast to this analysis, our main analyses consider all systematic AI signals that are not due to CNA, and find its contribution to be substantial and comparable to the CNA.

Additionally, for the non-CNA mechanisms, their quantification is likely to be noisier (e.g. estimating the effects of non-coding variants is notoriously difficult), while large CNA can be estimated fairly accurately, further biasing the apparent effect in favor of CNA-AI in this type of analysis that quantifies explainable variance. In contrast, in our analyses we quantify all systematic AI, regardless of the source, and by a subtraction of the CNA-attributable AI arrive at the 46.6% estimate of AI from non-CNA sources.

We note, however, that our reg-AI signal captures a composite of regulatory influences, alongside residual expression from contaminating normal cells (which we attempted to minimize through our modeling) and potential statistical artefacts. Similarly, we note that our CNA-AI estimate will also include effects of subclonal mutations, since they register as an imbalance at the DNA level similarly as the CNA do. However, our analysis demonstrates that subclonal mutations account for only a minor fraction of significant CNA-AI calls in cancer genes (Figure S2). Additionally, subclonality can only generate negative CNA-AI, while our subsequent analyses focus on positive AI signals from any source (CNA, regulatory, or overall mRNA-AI).

*Allele-specific chromatin accessibility (ASCA) analysis*

We performed variant calling on ATAC-Seq data using the same pipeline applied to RNA-Seq data to quantify allele-specific chromatin accessibility (ASCA). Variants were filtered to include only those with at least 5 ATAC-Seq read counts and a “PASS” at the FILTER column obtained from WES variant calling.

We applied the beta-binomial modeling framework described in "Beta-binomial modeling of allelic imbalance" to quantify chromatin accessibility imbalances corrected for copy number alterations, substituting ATAC-Seq read counts for RNA-Seq counts in the model. This approach allowed us to calculate copy number-corrected ASCA analogous to our reg-AI calculations.

For correlation analysis between ASCA and reg-AI, we selected matched variants from WES, RNA-Seq and ATAC-Seq for our analysis, and kept those having more than 20 total (i.e. sum of WT and MUT alleles) RNA-Seq and ATAC-Seq reads (n = 782). We computed Pearson product-moment correlation coefficients (R) between the residuals from our copy number-corrected models for both datasets: $\Delta ATAC\_VAF(ATAC\_VAF_{obs} - ATAC\_VAF_{pred})$ from ATAC-Seq (ASCA residuals) versus $\Delta RNA\_VAF(RNA\_VAF_{obs} - RNA\_VAF_{pred})$ from RNA-Seq (reg-AI residuals).

*Gene-level DNA methylation status*

We sourced DNA methylation data from the TCGA, represented as a matrix of beta-values for ~450k CpG sites across all samples and cancers. Initial processing involved several filtering steps: (1) Removal of cross-reactive CpGs[3,4]; (2) exclusion of probes with missing values (NAs) in more than 20% of samples, applied separately for each cancer type; (3) when multiple samples existed for an individual, average methylation values were computed.

Before analysis, beta-values were converted to M-values using the *beta2m* function from the *lumi* R package. We focused on CpGs near the TSS to examine regions likely influencing transcription initiation. Specifically, we averaged M-values across CpGs located within "TSS1500", "TSS200", "1stExon", and "5’ UTR" categories per the Illumina 450k Manifest (https://support.illumina.com/downloads/infinium_humanmethylation450_product_files.html). We included additional probes within the first 2kb of the gene body as well, determined by distances from the TSS to CpGs using gene annotations from GENCODE v22 (https://zwdzwd.github.io/InfiniumAnnotation). Only CpGs within a 2kb range from the main isoform's TSS (using MANE as a reference isoform[5]), were included.

This approach yielded a single M-value per gene per individual, reflecting gene-specific DNA methylation levels near the promoter region. We then categorized this gene promoter-proximal DNA methylation status across the entire cohort based on percentile ranks separately per gene: samples with M-values in the lowest 30% for a given gene were labeled as unmethylated, those above the 70% percentile as methylated, and those in between as partially methylated (as would result from one allele being methylated and the other unmethylated). These methylation statuses were subsequently correlated with our reg-AI values to explore potential relationships between methylation patterns and allele-specific expression.

*Variant effect predictions on chromatin profiles by Sei*

To predict the impact of coding mutations on chromatin state, we utilized the *Sei* tool[6], which uses deep neural networks to assess the chromatin-altering effects of genetic variants, within our AI dataset, with default parameters. *Sei* analyzes an extensive range of publicly available chromatin profiles—totaling 21,907—covering a wide spectrum of *cis*-regulatory elements, including transcription factor binding profiles, histone modification marks, and chromatin accessibility profiles, providing insights with single-nucleotide resolution. For simplicity, we employed the simplified categorical classification by *Sei*, focusing on four distinct "sequence classes", associated with active transcription (TN1-TN4). For each somatic mutation, we calculated the absolute maximum score across all four transcriptional states. Mutations were then classified based on whether their *Sei* scores exceeded the 95th percentile threshold, categorizing them as having either a predicted impact on transcription-associated chromatin states ("High-Impact") or remaining neutral.

*Variant effect predictions on transcription initiation by Puffin-D*

We estimated the transcriptional impact of exonic mutations using *Puffin-D*[7], a deep learning model designed to predict transcription initiation signals across a 100 kb DNA sequence. This model allows us to assess promoter activity based on experimental data sources, including GRO/PRO-Seq, FANTOM, and ENCODE CAGE scores.

For our analysis, we focused on 100 kb-wide windows centered around each mutation ([-50 kb, +50 kb]). Both the *wild-type* and mutated DNA sequences were used as inputs (one sequence per mutation). *Puffin-D* generates a score for each nucleotide within this sequence, allowing us to measure the transcriptional impact of mutations. We calculated the delta *Puffin-D* score (Δ*Puffin-D*) as the difference between *wild-type* and mutated sequences for each nucleotide position.

To focus on transcription initiation effects, we analyzed a ±50 bp window centered on the gene's main TSS, as defined by the MANE Select transcripts from GENCODE v26. We summed the absolute Δ*Puffin-D* scores within this narrow window yielding a single score per somatic mutation in our dataset for each of the three prediction datasets (GRO-Seq, PRO-Seq, and FANTOM CAGE), then selected the maximum score across all three predictions as the final Δ*Puffin-D* score (ENCODE excluded).

To interpret the magnitude of Δ*Puffin-D* scores, we used established positive controls. These controls included known promoter mutations in the oncogene *TERT* with known effects on transcriptional activity[8,9]. We created artificial sequences with these promoter mutations and compared their *Puffin-D* scores to those of the corresponding *wild-type* sequences, as stated above. After calculating the single Δ*Puffin-D* score for each positive control mutation, we compared these values to the Δ*Puffin-D* scores of our somatic mutations to benchmark their potential impact on transcription initiation.

*Variant effect predictions on splicing and enrichment analysis*

To evaluate the impact of splicing alterations on AI in our dataset (792,303 unique somatic mutations), we utilized two deep learning-based tools: *SpliceAI*[10] and *Pangolin*[11]. Both tools employ deep residual neural networks to predict splicing changes at each position within a 50-bp window of a pre-mRNA transcript based on its genomic sequence.

*SpliceAI* delta scores quantify changes in splice site activity due to mutations, across four categories: Donor Loss ($SP_{DL}$), Donor Gain ($SP_{DG}$), Acceptor Loss ($SP_{AL}$), and Acceptor Gain ($SP_{AG}$). These delta scores reflect the increase or decrease in splice site usage between the mutated and reference alleles. To capture the most impactful potential splicing alteration for each variant, we defined an aggregate score, the $SP_{Max\Delta}$, as the maximum absolute delta score across these four categories. Mutations with *SpliceAI* delta scores ≥ 0.15 were classified as "splice-disrupting", while others were considered "splice-neutral". This threshold was adapted from the original *SpliceAI* publication, which suggested a threshold of 0.2; we selected a slightly more relaxed threshold of 0.15 to increase the number of variants considered for potential splice effects in our analysis.

*Pangolin* reports two analogous delta scores representing the maximum predicted increase ($PG_{gain}$) or decrease ($PG_{loss}$) in splice site activity. Similarly to our approach with *SpliceAI*, we defined a $PG_{Max\Delta}$ score as the maximum absolute value of these two *Pangolin* scores (i.e., max($|PG_{gain}|$, $|PG_{loss}|$)). Mutations with *Pangolin* delta scores ≥ 0.10 were classified as "splice-disrupting", while others were considered "splice-neutral". As the original *Pangolin* manuscript did not specify a definitive threshold for splice disruption, we established this 0.10 threshold by correlating $PG_{Max\Delta}$ scores with $SP_{Max\Delta}$ scores across our dataset (Pearson correlation coefficient, r = 0.75). A Locally Estimated Scatterplot Smoothing (LOESS) regression fitted between $SP_{Max\Delta}$ and $PG_{Max\Delta}$ indicated that a SpliceAI $SP_{Max\Delta}$ score of 0.15 corresponded approximately to a *Pangolin* $PG_{Max\Delta}$ score of 0.10.

Following the classification of SNVs based on their predicted splicing effects, we performed enrichment analysis to determine whether the proportion of predicted splice-disrupting variants was significantly higher for mutations associated with negative reg-AI (and separately for positive reg-AI) compared to those with no significant reg-AI. These analyses were performed for distinct gene category (OGs, TSGs, or passenger genes as controls) and predicted functional impact of SNVs (high aa-impact missense or effectively synonymous; also stratified further into low aa-impact missense and synonymous-only). Enrichment analyses were performed for each of the five *SpliceAI* delta score categories, and three Pangolin delta score categories, calculating ORs and *p*-values using Fisher's Exact test.
This enrichment analysis was conducted at two levels:

Gene category-level enrichment: Applied across gene categories including cognate and noncognate OGs, cognate and noncognate TSGs, *TP53* as a separate category, and passenger genes.

Individual gene-level enrichment: Focused on the same subset of known cancer driver genes used in our AI-dN/dES tests. Following the enrichment analyses, FDR corrections were applied separately for each stratification—by SNV type, *SpliceAI/Pangolin* delta score category used, and reg-AI direction (positive or negative). Genes were considered significant where FDR < 10%.

*<u>Classification of nonsense mutations into NMD-triggering and NMD-evading</u>*

Nonsense mutations often lead to nonsense-mediated mRNA decay (NMD) by creating premature termination codons (PTCs) on the coding sequence. However, some PTCs can evade this pathway, allowing mRNA to escape degradation. We classified nonsense mutations as NMD-triggering or NMD-evading based on established genomic rules[12]: (1) *55nt-rule:* nonsense mutations located ≤ 55 nt downstream from the last base of the penultimate exon are categorized as NMD-evading; (2) *Last exon rule:* Nonsense mutations within the last exon are considered NMD-evading; (3) *Start-proximal rule:* Nonsense mutations within the first 200 nt of the transcript are deemed NMD-evading.
